# Isoform-specific roles of the *Drosophila* filamin-type protein Jitterbug (Jbug) during development

**DOI:** 10.1101/2020.10.13.337626

**Authors:** SeYeon Chung, Thao Phuong Le, Vishakha Vishwakarma, Yim Ling Cheng, Deborah J. Andrew

**Affiliations:** Department of Biological Sciences, Louisiana State University, Baton Rouge, LA 70803, USA; Department of Cell Biology, Johns Hopkins University School of Medicine, Baltimore, MD 21205, USA

**Author notes:** These authors contributed equally to this work (SeYeon Chung, Thao Phuong Le).

**Keywords:** *Drosophila*, filamin, *jitterbug*, actin, thoracic bristles, epidermal denticles, tubular epithelial organs, trans-splicing, Crb

## Abstract

Filamins are highly conserved actin-crosslinking proteins that regulate organization of the actin cytoskeleton. As key components of versatile signaling scaffolds, filamins are implicated in developmental anomalies and cancer. Multiple isoforms of filamins exist, raising the possibility of distinct functions for each isoform during development and in disease. Here, we provide an initial characterization of *jitterbug* (*jbug*), which encodes one of the two filamin-type proteins in *Drosophila*. We generate Jbug antiserum that recognizes all of the spliced forms and reveals differential expression of different Jbug isoforms during development, and a significant maternal contribution of Jbug protein. To reveal the function of Jbug isoforms, we create new genetic tools, including a null allele that deletes all isoforms, hypomorphic alleles that affect only a subset, and UAS lines for Gal4-driven expression of the major isoforms. Using these tools, we demonstrate that Jbug is required for viability and that specific isoforms are required in the formation of actin-rich protrusions including thoracic bristles in adults and ventral denticles in the embryo. We also show that specific isoforms of Jbug show differential localization within epithelia and that maternal and zygotic loss of *jbug* disrupts Crumbs (Crb) localization in several epithelial cell types.

## Introduction

Actin-crosslinking proteins organize actin filaments into higher-order structures such as orthogonal actin arrays and parallel actin bundles. Filamins are highly conserved large cytoplasmic proteins that crosslink actin filaments into dynamic three-dimensional structures (Nakamura *et al*. 2011; Razinia *et al*. 2012). Filamins consist of actin-binding Calponin-homology (CH) domains at their N-terminus, followed by a long C-terminal rod-like domain of filamin-type immunoglobulin (IG-FLMN) repeats. Dimerization of filamins through the last C-terminal IG-FLMN repeat allows the formation of a flexible V-shaped structure that holds two actin filaments at large angles to create either a loose three-dimensional actin network (Nakamura *et al*. 2011; Razinia *et al*. 2012) or parallel bundles of actin filaments (Sokol and Cooley 1999; Gay *et al*. 2011). In addition to binding to actin, filamins interact with transmembrane receptors, adhesion molecules and even transcription factors, and are thus involved in multiple cell functions, including motility, maintenance of cell shape, and differentiation (Nakamura *et al*. 2011; Razinia *et al*. 2012). Mutations in filamins are associated with a wide range of congenital anomalies and have been shown to both promote and inhibit metastasis and cancer growth (Razinia *et al*. 2012; Savoy and Ghosh 2013; Shao *et al*. 2016; Sasaki *et al*. 2019), emphasizing the multiple critical roles of filamins in development and disease.

Studies in *Drosophila* have helped refine our understanding of the roles of filamins in vivo. *Drosophila* has two orthologs of human filamin A: Cheerio (Cher) and Jitterbug (Jbug). Cher shares the organization of the protein and 46% identity and 61% similarity in amino acid sequence with human filamin A. Jbug, with 23% identity and 36% similarity in amino acid sequence with human filamin A, has some distinct features, including an actin-binding domain consisting of three, instead of two, CH domains. Several studies have revealed roles for Cher in organizing the F-actin cytoskeleton in multiple developmental contexts and diseases, including formation of ovarian germline ring canals (Li *et al*. 1999; Sokol and Cooley 1999; Huelsmann *et al*. 2016), follicle cell migration (Sokol and Cooley 2003), tissue morphogenesis during cellularization (Krueger *et al*. 2019), and *Drosophila* tumorigenesis (Külshammer and Uhlirova 2013). Compared to the extensive studies on Cher, our understanding of the roles of Jbug is more limited. Originally identified as a gene that causes bang-sensitive seizure – the phenotype that gives the gene its *jitterbug* name (Song and Tanouye 2006), *jbug* is required in photoreceptor cells for axon targeting (Oliva *et al*. 2015) and in tendon cells to maintain their shape at the muscle-tendon junction (Olguín *et al*. 2011; Manieu *et al*. 2018).

Multiple isoforms of filamin proteins exist in both *Drosophila* and mammals (Sokol and Cooley 1999; Browne *et al*. 2000; Van der Flier and Sonnenberg 2001; Gorlin *et al*. 1990; Wang *et al*. 2007). Originally, only two Cher isoforms were reported: a large isoform (FLN240) that contains both actin-binding CH domains and IG-FLMN repeats and a smaller one (FLN90) that contains only IG-FLMN repeats (Sokol and Cooley 1999). Recent annotations indicate that *cher* encodes six longer isoforms (>240 kDa), which include two actin-binding CH domains and 18-22 IG-FLMN repeats, and four shorter isoforms (90-100 kDa), which include only eight IG-FLMN repeats without any CH domains (Flybase; www.flybase.org; Thurmond *et al*. 2019). Importantly, a recent study in *Drosophila* larval neuromuscular junctions revealed a role for the shorter Cher isoform (FLN90) as a postsynaptic scaffold in *Drosophila* larval neuromuscular junctions during synapse formation (Lee and Schwarz 2016), suggesting distinct roles for different isoforms. Like *cher*, *jbug* also encodes multiple isoforms with a range of protein sizes. However, the roles for *jbug* have been revealed mostly using RNA interference (RNAi) constructs that target most of the existing splice forms (Olguín *et al*. 2011; Manieu *et al*. 2018), preventing the characterization of isoform-specific roles for Jbug.

Here, we report on the creation of several new genetic tools for parsing out the roles of the different Jbug isoforms, including a null allele that deletes all isoforms and hypomorphic alleles that affect only a few. We also create UAS constructs for expressing the major Jbug isoforms. We demonstrate that different isoforms are differentially expressed during development and localize to different domains in embryonic epithelia. We show isoform-specific roles of Jbug in formation of epidermal denticles during embryogenesis and thoracic bristles in adults. Finally, we also report that removal of both maternal and tissue-specific supplies of *jbug* disrupts tissue morphology and reduces Crb accumulation in the subapical domain of epithelial cells.

## Materials and Methods

### Fly stocks and husbandry

Fly lines used in our experiments were: *Oregon R*; *jbug^20^*, *jbug^30^*, *jbug^133^*, UAS-Jbug-RC, UAS-Jbug-RI, UAS-Jbug-RF (this work); UAS-Jbug-RL (Olguín *et al*. 2011); *fkh-Gal4* (Henderson and Andrew 2000); *69B-Gal4* (Rorth 1996); *da-Gal4* (Wodarz *et al*. 1995); *matα-Gal4* (Häcker and Perrimon 1998), *Df(2R)59AB*, *Df(2R)Exel6079*, *TRiP.JF01166* (Bloomington Stock Center); *GD8664*, *GD13033* (Vienna *Drosophila* RNAi Center). All crosses were performed at 25°C.

### Generation of *jbug* mutant alleles

*jbug* mutants were generated by homologous recombination (Gong and Golic 2003). For *jbug^20^* null mutants, genomic fragments upstream and downstream of the *jbug* ORF were amplified by PCR using the following primers (restriction sites in bold and linkers in italic):

in-fusion_L-22737893, 5’-*CTAGTCTAGGGCGCGCC*GATGAGTTGTGGCTTGAGCA-3’

in-fusion_R-22742442, 5’-*TAGGGGATCACGTACG*GCCAAACCAATGCTGAAGAT-3’

NotI_L-22717574, 5’-AT**GCGGCCGC**TAGAGGGGAGAGCAAGTGGA-3’

Acc65I_R-22721186, 5’-ATA**GGTACC**GTGTGAGCTTCGGGATCAAT-3’

For *jbug^30^* and *jbug^133^* hypomorphic alleles, genomic fragments upstream and downstream of an exon common to all *jbug* splice forms were amplified using the following primers (restriction sites in bold):

AscI_L-22727168, 5’-AT**GGCGCGCC**AAAGCCGTTGAAGAAAGCAA-3’

BsiWI_R-22731840, 5’-ATA**CGTACG**CTCGATGCACTTTGTCTCCA-3’

NotI_L-22717574, 5’-AT**GCGGCCGC**TAGAGGGGAGAGCAAGTGGA-3’

Acc65I_R-22721186, 5’-ATA**GGTACC**GTGTGAGCTTCGGGATCAAT-3’

PCR fragments were cloned into pW25, which carries *white^+^*, the recognition site for I-SceI endonuclease, and FRT sites. The constructs were injected into *w^1118^* embryos by Rainbow Transgenic Flies, Inc. Transformants were crossed to flies carrying hs-I-SceI and hs-Flp, and progeny were heat shocked (37°C) for 1 hour 48–72 hours after egg laying (AEL). Recombination and insertion were confirmed by genomic PCR and reverse-transcriptase PCR (RT-PCR) followed by sequencing.

The following primers were used for RT-PCR for *jbug^20^* shown in Supplemental Figure 2 (restriction sites in bold and linkers in italic):

*jbug* 5-1, 5’-GGAGAACAAGTACCGGGTGA-3’

*jbug* 5-2, 5’-*GGATCCTCGAGCATATG*TCCTTCCACGTTCACCTCGA-3’ (same as Jbug-Ab-3’ for making Jbug antibody)

*jbug* 3-1, 5’-AT**GCGGCCGC**TAGAGGGGAGAGCAAGTGGA-3’ (same as NotI_L-22717574 used for amplifying a homology arm downstream of the *jbug* ORF)

*jbug* 3-2, 5’-*CGCGCGGCAGCCATATG*ATGTCCTCACCCGGCCTAAC-3’ (same as Jbug-Ab-5’ for making Jbug antibody)

### Generation of UAS-Jbug-RC, -RI and -RF transgenic flies

An ORF for *jbug-RC* (1230 bp) was PCR-amplified using the existing cDNA RE40504 as a template. A full length cDNA for *jbug-RF* (9430 bp) was created by In-Fusion cloning of three fragments. The 5’ fragment was prepared by enzyme digestion of RE40504 (cut with ApaI and BamHI). The middle fragment (1189 bp) was amplified by reverse transcriptase-PCR (RT-PCR), and the ApaI and BglII sites were added to the 5’ and the 3’ end of the fragment, respectively. The 3’ fragment was amplified by PCR using the existing cDNA GH03118 from the BglII site to the end of the ORF, and the BamHI site was added to the 3’ end. The ligated full cDNA was confirmed by sequencing. A full cDNA for *jbug-RI* (5989 bp) was created by PCR amplification using the *jbug-RF* cDNA as a template and confirmed by sequencing. *jbug-RC*, *jbug-RI* and *jbug-RF* ORFs were PCR-amplified and cloned into the pUAST vector (Brand and Perrimon, 1993) to create UAS lines. The DNA constructs were injected into *w^1118^* embryos by Rainbow Transgenics, Inc. (UAS-Jbug-RC) and GenetiVision (UAS-Jbug-RI and UAS-Jbug-RF).

### Generation of Jbug antibody

A PCR fragment of 1212 bp common to all *jbug* splice forms was cloned into the pET15b vector. The fragment corresponds to almost the entire open reading frame for the Jbug-PC isoform except for the last six amino acids and contains four IG-FLMN repeats. Primers used for In-Fusion cloning were (linkers in italic):

Jbug-Ab-5’, 5’- *CGCGCGGCAGCCATATG*ATGTCCTCACCCGGCCTAAC-3’

Jbug-Ab-3’, 5’- *GGATCCTCGAGCATATG*TCCTTCCACGTTCACCTCGA-3’

The construct was transformed into BL21-DE3 cells for protein induction with IPTG. Recombinant protein was purified using Ni-NTA agarose (Qiagen). Rabbit polyclonal antibodies were generated by Covance, Inc. and used at a dilution of 1:500 for immunohistochemistry and 1:5,000 for western blot.

### Whole-mount in situ hybridization

In situ hybridization was performed as described previously (Lehmann and Tautz 1994). Probes specific for a subset of *jbug* splice forms are shown in Figure 1A. DNA fragments for probes 1 and 3 were PCR-amplified using wild-type genomic DNA, cloned into the pCR2.1 vector and sequenced to verify. Probe 2 was made from cDNA GH03013, linearized at the KpnI site to create a probe specific to all jbug splice forms. Antisense digoxigenin-labeled RNA probes were generated using T7 (probes 1 and 3) or SP6 (probe 2) polymerase on linearized templates. The molecular regions corresponding to the RNA probes are as follows:

**Figure 1.**
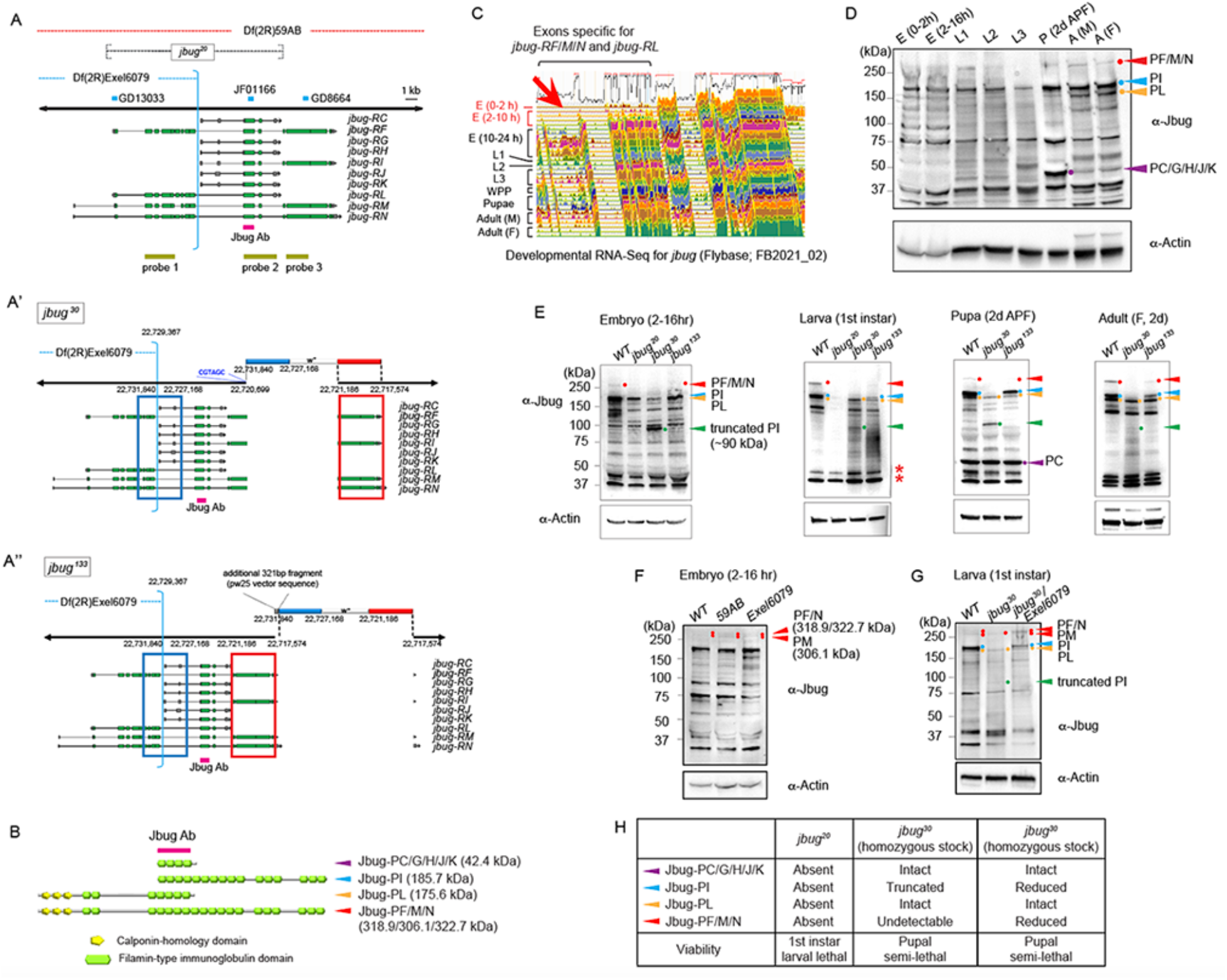
Differential expression of Jbug isoforms throughout *Drosophila* development and generation of null and hypomorphic alleles of *jbug*. (A) Ten annotated splice forms of *jbug* are reported by FlyBase. Deletion regions for two deficiency lines in the *jbug* locus and the *jbug^20^* null allele are indicated. Cyan bars, three RNAi lines for *jbug* that were tested in this study. Magenta bar, a common exon for all isoforms that was used to generate an antibody against Jbug. Light green bars, regions used for making antisense probes for in situ hybridization. (A’, A’’) Molecular alterations in two hypomorphic alleles of *jbug*. Blue and red boxes represent homology arms that were used for homologous recombination. Genomic sequence locations are shown for the homology arms. Hypomorphic alleles were generated by incomplete homologous recombination, with a partial (*jbug^30^*, A’) or a complete (*jbug^133^*, A’’) insertion of the donor DNA into the *jbug* isoforms that have a longer C terminus (*RF/M/N* and *RI*). (B) Six different Jbug protein isoforms are made from the different annotated splice forms. (C) Developmental RNA-Seq data by Flybase (www.flybase.org; FB2021_02). Note that only minimal expression of *jbug-RF*/*M/N* and *jbug-RL* is detected until 10 hours of embryogenesis (red arrow). (D-F) Western blot using protein extracts from wild type and *jbug* mutants at different stages. Bands that correspond to the predicted molecular weights of several isoforms are marked with dots and arrowheads with different colors. Arrowheads of the same colors are used to indicate the major isoforms in B. (D) Developmental western blot using protein extracts from embryos (0-2 and 2-16 hours AEL at 25°C), 1st, 2nd and 3rd instar larvae, pupae (2 days APF) and male (M) and female (F) adults (2 days post-eclosion). (E) Western blots using protein extracts from different *jbug* mutants. Asterisks, two small bands that are still observed in *jbug^20^* larvae, which are likely from nonspecific cross-reactivity. (F) In both *59AB* and *Exel6079* homozygous embryos, the same protein bands are detected as in wild type. Note that two faint but distinct bands are visible for >300 KDa, suggesting that Jbug-PM (306.1 kDa) and the other two bigger isoforms, Jbug-PF (318.9 kDa) and Jbug-PN (322.7 kDa), are separated in this gel. (J) In *jbug^30^/Exel6079* transheterozygous larvae, Jbug-PF/M/N, Jbug-PI and Jbug-PL bands are all detected. (H) Protein levels of Jbug isoforms in each *jbug* mutant. Each mutant’s viability is also shown.

Probe 1 (specific for *jbug-RF/M/N* and *jbug-RL*): 22734199 – 22731666

Probe 2 (detects all *jbug* transcripts): 22722586-22722906, 22723936-22724162, 22724531-22725474

Probe 3 (specific for *jbug-RF/M/N* and *jbug-RI*): 22721775 – 22720075

### Western blot

Protein extracts from embryos 0-2 and 2-16 hours AEL, 1^st^, 2^nd^, 3^rd^ instar larvae, pupae (2 days after puparium formation (APF), and 2-day old male and female adults were used. For embryos homozygous for *jbug^20^*, *Df(2R)59AB* and *Df(2R)Exel6079* and for 1^st^ instar larvae homozygous for *jbug^20^* and transheterozygous for *jbug^30^*/*Df(2R)Exel6079*, a balancer chromosome that contains the twi-Gal4 UAS-GFP construct was used to distinguish desired genotypes from heterozygous siblings. Rabbit *α*-Jbug (this work; 1:5,000), mouse *α*-Actin (RRID:AB_11004139; Invitrogen; 1:2,000), and HRP-labelled secondary antibodies (RRID:AB_430833; RRID:AB_430834; Promega; 1:2,000) were used. Protein bands were detected using Pierce ECL Western Blotting Substrate.

### Scanning electron microscopy

Wild type (*Oregon R*) and *jbug^30^* homozygous male flies were coated with gold palladium and examined and photographed in a LEO/Zeiss Field-emission SEM.

### Antibody staining and confocal microscopy

Embryos were collected on grape juice-agar plates and processed for immunofluorescence using standard procedures. Briefly, embryos were dechorionated in 50% bleach, fixed in 1:1 heptane:formaldehyde for 40 min and devitellinized with 80% EtOH, then stained with primary and secondary antibodies in PBSTB (1X PBS, 0.1% Triton X-100, 0.2% BSA). For Jbug and phalloidin staining, embryos were hand-devitellinized. Antibodies used include: mouse *α*-Crb (RRID:AB_528181; DSHB (Cq4); 1:10), mouse α-α-Spec (RRID:AB_528473; DSHB (3A9); 1:2), rabbit α-Neurexin IV (Baumgartner *et al*. 1996; 1:2,000), rabbit α-CrebA (RRID:AB_10805295; 1:5,000), guinea pig α-Sage (Fox *et al*. 2013; 1:100), chicken α-GFP (RRID:AB_2534023; Invitrogen; 1:500), rat α-E-Cad (RRID:AB_528120; DSHB (DCAD2); 1:50), rabbit α-Jbug (this work; 1:500), rabbit α-β-Gal (RRID:AB_221539; Invitrogen; 1:500). Alexa fluor 488, 568, 647-labelled secondary antibodies were used at 1:500 (RRID:AB_142924; RRID:AB_141778; RRID:AB_2535766; RRID:AB_143157; RRID:AB_2535812; Molecular Probes). Alexa fluor-568-labeled phalloidin was used for F-actin labelling (RRID:AB_2632953; Molecular Probes; 1:250). All images were taken with a Leica SP8 confocal microscope using either a 63x, 1.4 NA or a 40x, 1.3 NA oil objective and the LAS X software (Leica).

### Quantification of denticle precursors in the ventral epidermis

Embryos stained for phalloidin and E-Cad were imaged with a Leica SP8 confocal microscope. A maximum intensity projection was generated using two focal planes (0.3 μm apart) in the region of adherens junctions. Denticle belts in abdominal segments 4-6 in stage 16 embryos were selected for quantification. Six columns of cells from anterior to posterior of each belt were analyzed. In each column, denticle precursors of four cells closest to the ventral midline were counted. For each genotype, five to eight embryos were quantified. Unpaired Student’s t-test with Welch’s correction was used for statistical analysis.

### Data availability

All data and methods required to confirm the conclusions of this work are within the article, figures, and supplemental materials.

## Results

### Different splice forms of *jbug* are differentially expressed during development

Ten annotated splice forms of *jbug*, which encode six different protein isoforms, are reported in Flybase (www.flybase.org; Thurmond *et al*. 2019). All isoforms contain multiple repeats of filamin-type immunoglobulin (IG-FMLN) domains (Figure 1, A and B), which form a rod-like structure in filamin. Two Jbug isoforms consist exclusively of these repeats; Jbug-PC/G/H/J/K (each splice form encodes the same 42.4 kDa protein; hereafter referred to as Jbug-PC) has four IG-FMLN repeats and Jbug-PI (185.7 kDa) has sixteen (Figure 1B). On the other hand, Jbug-PL (175.6 kDa) and Jbug-PF/M/N, the three longest Jbug isoforms (>300 kDa), contain three tandem calponin-homology (CH) actin-binding domains in their N-termini as well as multiple IG-FMNLs in their C-termini (seven and nineteen IG-FMLNs for Jbug-PL and Jbug-PF/M/N, respectively) (Figure 1B).

High-throughput expression data available in Flybase (www.flybase.org; Thurmond *et al*. 2019) showed high expression of *jbug* splice forms throughout all developmental stages, except during 0-10 hours of embryogenesis when there is a minimal expression of *jbug-RF/M/N* and *jbug-RL* (Figure 1C). BDGP Expression Data showed that, in addition to the strong ectodermal expression at earlier stages, *jbug* is also upregulated in epithelial tubular organs, including the trachea primordia and the salivary gland (https://insitu.fruitfly.org/cgi-bin/ex/insitu.pl). Our previous work confirmed this expression and revealed that *jbug* tracheal expression requires Trachealess, a major transcription factor in the trachea (Chung *et al*. 2011).

in situ hybridization using probes specific for different *jbug* splice forms further revealed tissue-specific expression of specific *jbug* splice forms (Supplemental Figure 1). A 5’ probe (Figure 1A; probe1) that should detect *jbug-RL* (the form that includes the CH actin-binding domains) and the longest splice forms *jbug-RF/M/N* revealed no expression until embryonic stage 12, when expression in the clypeolabrum (CL) was first detected at low levels (Supplemental Figure 1, left column). This CL expression persisted through embryogenesis with later additional expression in another group of anterior cells that, together with the CL-positive cells, forms the mouth and low level expression in posterior cells that form the posterior spiracles. A 3’ probe (Figure 1A; probe 3) that detects *jbug-RI* (the form that includes sixteen IG-FLMN repeats without the CH actin-binding domains) and *jbug-RF/M/N* reveals broad ectodermal expression in early embryos, with tissue-specific expression apparent by stage 11 in the CL, salivary gland, and tracheal pits (tp), with some low level expression in the foregut and subset of midline cells (Supplemental Figure 1; right column). By stage 12, the CL and tp staining are no longer detectable and expression is now also observed in epidermal stripes. By late stages, expression is also observed in the region of the posterior spiracles. A probe that detects all *jbug* splice forms (Figure 1A, probe 2) revealed a pattern that was the sum of the expression patterns observed with the more 5’ and 3’ isoform-specific probes. Given the very limited overlap in staining patterns with the 5’ (probe 1) and 3’ (probe 3) probes, this finding suggests that *jbug-RL* and *jbug-RI* are the major mRNA splice forms made in embryos and that *jbug-RL* is much more restricted in its expression than *jbug-RI*, both in terms of how early the transcripts are detected and how broadly expression is observed. If *jbug RF/M/N* transcripts are produced in embryos, the expression must be limited to the late posterior spiracles, the only site where all three probes detected signals. Likewise, if *jbug-RC* is made in embryos, it must completely overlap the expression patterns of *jbug-RL* and *jbug-RI*.

To analyze protein expression levels of Jbug during *Drosophila* development, we generated Jbug antiserum that recognizes a protein domain encoded by all ten *jbug* splice forms (Figure 1A). This domain includes the four IG-FLMN repeats present in all Jbug isoforms (Figure 1B). Our Western blot analysis using wild-type protein extracts revealed that Jbug proteins are detected at high levels throughout all developmental stages (Figure 1D). Strong Jbug signals in protein extracts from 0-2 hour embryos (Figure 1D) also suggest a maternally provided pool of Jbug. The Jbug antiserum detected bands in size ranges predicted from the major splice forms (42.4 kDa, 175.6 kDa, 185.7 kDa, >300 kDa). Importantly, the different isoforms were differentially expressed during development (Figure 1D). Jbug-PC (42.4 KDa) was abundant only in pupae and Jbug-PL (175.6 kDa) was most abundant in adults (Figure 1D). Bands corresponding in size to Jbug-PI (185.7 kDa) and Jbug-PF/M/N (>300 kDa) were detected at all stages with slight differences in protein levels. The PI band had the highest intensity at almost all stages, with the exception of PC in pupae, where the levels of PI and PC were similar (Figure 1D). Jbug-PF/M/N isoforms differ only slightly from one another in regions outside the CH and IG-FLMN domains. Due to their huge sizes and a very slight difference in molecular weight among Jbug-PF/M/N, we were not able to distinguish exactly which of the three isoforms is expressed at any specific stage. The large size of these isoforms potentially also diminishes membrane transfer, potentially leading to relatively weak signals. Alternatively, the relatively weak signals could indicate that only very low levels of the longest splice forms are made. Notably, several intermediate-sized bands that did not match the predicted size of annotated isoforms were also detected at different levels throughout development (Figure 1D). The existence of these bands suggests either that more Jbug isoforms exist than have been annotated or that proteolyzed and/or phosphorylated fragments are being detected (also see Figure 1E; most of these extra bands are absent in *jbug* null mutant larval extracts). Indeed, roles for proteolytic forms of Filamin A (Bedolla *et al*. 2009; Savoy and Ghosh 2013; Yue *et al*. 2013) as well as phosphorylated Filamin A (Ohta and Hartwig 1996; Tirupula *et al*. 2015) have been reported from studies of the mammalian proteins. Overall, our data indicate that different splice forms of *jbug* are differentially expressed during *Drosophila* development.

### Null and hypomorphic *jbug* alleles have been generated

To investigate roles for *jbug* in development, we generated mutant alleles for *jbug* using homologous recombination (Gong and Golic 2003). Two donor DNA constructs were used to target different genomic regions. One construct targets the genomic region encompassing the translation start sites for all splice forms and extends through the exons shared by all splice forms; the other targets only the region containing the exons shared by all splice forms. With the construct that targets the larger region, we obtained a null mutant, *jbug^20^* (Figure 1A). PCR analysis confirmed that the entire targeted region was replaced with the mini-*white^+^* (mini-*w^+^*) gene (data not shown). Reverse transcriptase-PCR (RT-PCR) using mRNA extracted from homozygous *jbug^20^* embryos further confirmed that no *jbug* transcripts are produced in *jbug^20^* embryos (Supplemental Figure 2, A-C). Both *jbug^20^* homozygous animals and transheterozygotes for *jbug^20^* over either of the two deficiency lines that delete all or part of the *jbug* locus (*59AB* and *Exel6079*; Figure 1A) died during the first instar larval stage (Figure 1H; Table 1). Western blot analysis using Jbug antiserum showed that whereas a significant amount of the Jbug proteins were detected in *jbug^20^* embryos (Figure 1, E and H), almost all protein bands were absent by the first instar larval stage, with the exception of two low molecular weight bands that are likely to be from nonspecific cross-reactivity (Figure 1E). Similarly, the same Jbug isoforms were detected both in *59AB* and *Exel6079* homozygous embryos as observed in WT (Figure 1F). These data suggest that Jbug proteins from maternally provided *jbug* mRNA or protein persist through embryogenesis. Since only very low level *jbug* mRNA is detected in 0-2 hour wild-type embryos, based on RNA-Seq (Figure 1C, Flybase; www.flybase.org) and in situ hybridization (Fly-FISH; http://fly-fish.ccbr.utoronto.ca; BDGP; https://insitu.fruitfly.org/cgi-bin/ex/insitu.pl; this study), it is the Jbug protein and <u>not *jbug* mRNA</u>, that is maternally provided.

**Table 1.**
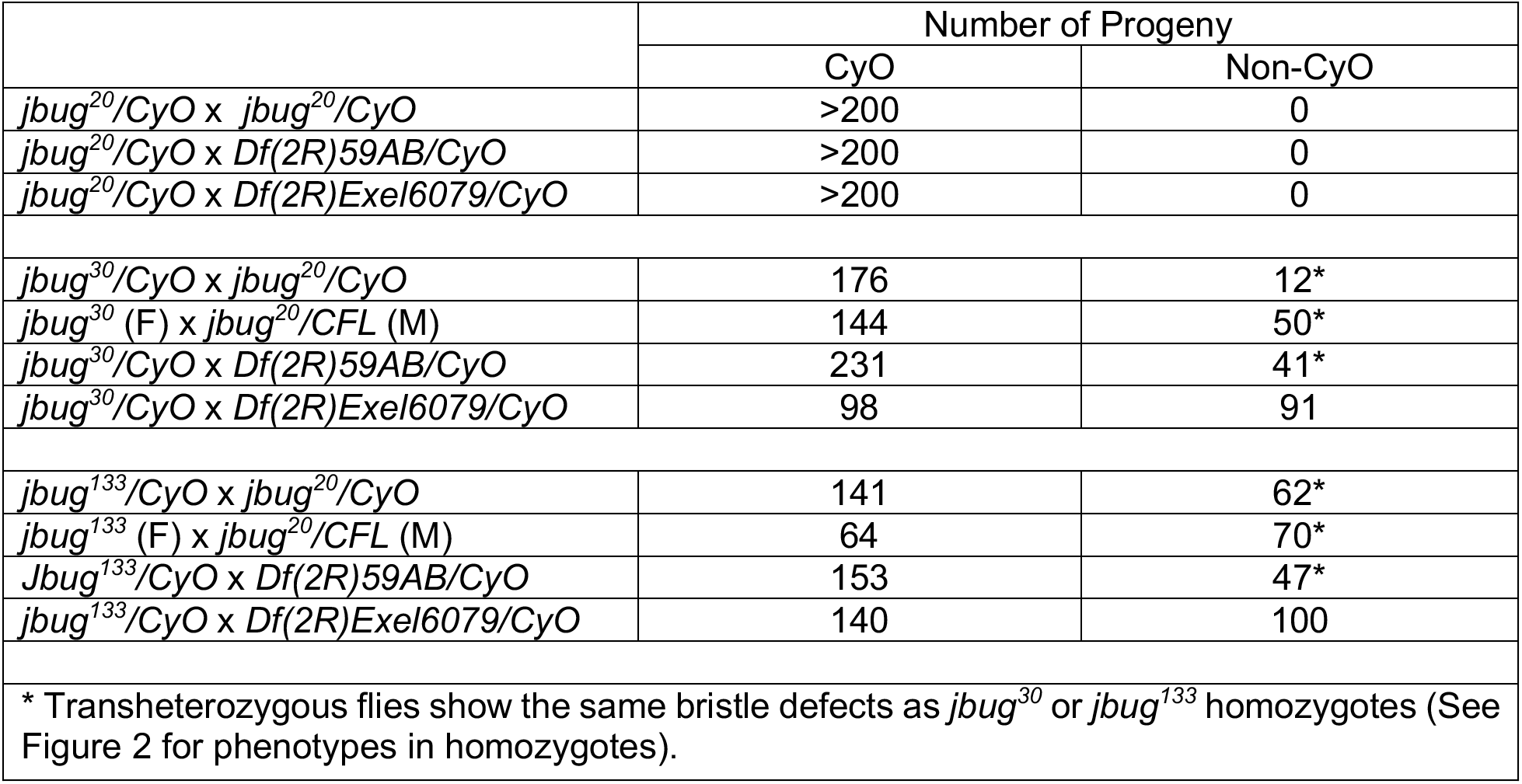
Complementation test.

|  | Number of Progeny |  |
| --- | --- | --- |
|  | CyO | Non-CyO |
| *jbug*<sup>20</sup>/CyO x *jbug*<sup>20</sup>/CyO | >200 | 0 |
| *jbug*<sup>20</sup>/CyO x *Df*(2R)59AB/CyO | >200 | 0 |
| *jbug*<sup>20</sup>/CyO x *Df*(2R)Exel6079/CyO | >200 | 0 |
| *jbug*<sup>30</sup>/CyO x *jbug*<sup>20</sup>/CyO | 176 | 12* |
| *jbug*<sup>30</sup> (F) x *jbug*<sup>20</sup>/CFL (M) | 144 | 50* |
| *jbug*<sup>30</sup>/CyO x *Df*(2R)59AB/CyO | 231 | 41* |
| *jbug*<sup>30</sup>/CyO x *Df*(2R)Exel6079/CyO | 98 | 91 |
| *jbug*<sup>133</sup>/CyO x *jbug*<sup>20</sup>/CyO | 141 | 62* |
| *jbug*<sup>133</sup> (F) x *jbug*<sup>20</sup>/CFL (M) | 64 | 70* |
| *Jbug*<sup>133</sup>/CyO x *Df*(2R)59AB/CyO | 153 | 47* |
| *jbug*<sup>133</sup>/CyO x *Df*(2R)Exel6079/CyO | 140 | 100 |
| * Transheterozygous flies show the same bristle defects as *jbug*<sup>30</sup> or *jbug*<sup>133</sup> homozygotes (See Figure 2 for phenotypes in homozygotes). |  |  |

With the DNA construct that aimed to remove the genomic region common for all splice forms of *jbug*, we unexpectedly obtained two hypomorphic alleles, *jbug^30^* and *jbug^133^*, that affect only a subset of splice forms. Incomplete recombination occurred in these two alleles, resulting in either a part of (for *jbug^30^*) or the entire DNA construct (for *jbug^133^*) being inserted into the *jbug* locus (Figure 1, A’ and A’’). In *jbug^30^*, the more 3’ homology arm in the donor DNA recombined correctly, but the other homology arm was inserted into the genome along with the mini*-w+* gene, rather than being recombined (Figure 1A’). In *jbug^133^*, homologous recombination failed to occur; instead, the entire donor construct was inserted into the 3’ UTR of the splice forms that have longer C-termini (*jbug-RI* and *jbug-RF*/*M*/*N*) (Figure 1A’’). These recombination events disrupted *jbug-RI* and *jbug-RF*/*M*/*N* in both alleles. Western blot analysis revealed that the Jbug-PF/M/N isoforms (>300 kDa) were either absent or decreased to undetectable levels in *jbug^30^* mutants at all stages (Figure 1E). Jbug-PI (185.7 kDa) was also absent in *jbug^30^*, and instead, a smaller protein (∼90 kDa) was detected in *jbug^30^* mutant animals at all stages (Figure 1, E and H). This additional band was not detected in wild-type or *jbug^133^* protein extracts at any stage, suggesting that insertion of the DNA construct generates a truncated form of Jbug-PI in *jbug^30^*. In *jbug^133^*, both Jbug-PF/M/N and Jbug-PI protein bands were slightly decreased in intensity (Figure 1, E and H), suggesting that insertion of the construct in the 3’ UTR region either affects mRNA stability or inhibits translation of these isoforms. Consistent with the molecular data, we did not detect any significant changes in the level of Jbug-PC (42.4 kDa) and Jbug-PL (175.6 kDa) in *jbug^30^* and *jbug^133^* protein extracts (Figure 1, E and H). Taken together, we generated new alleles for *jbug*, including a null mutant and two hypomorphic alleles that disrupt a subset of Jbug isoforms, specifically Jbug-PI and Jbug-PF/M/N.

### *jbug* hypomorphs show semi-lethality and defects in bristle development on the thorax

Many homozygous flies in *jbug^30^* and *jbug^133^* died during pupation, leaving a number of dead pupae that never eclosed. In both alleles, homozygous adult flies that survived were fertile and homozygous lines could be maintained. Interestingly, adult flies homozygous for *jbug^30^* or *jbug^133^* produced short, bent and rough-ended bristles on the thorax (Figure 2A), suggesting a role for Jbug in bristle development, perhaps in crosslinking actin bundles that form the bristles. Consistent with our findings that levels of Jbug-PF/M/N and Jbug-PI were more significantly reduced in *jbug^30^* than in *jbug^133^* (Figure 1, E and H), *jbug^30^* flies showed a higher expressivity of bristle defects than *jbug^133^*. Whereas almost every thoracic bristle had defects in *jbug^30^* homozygous adults, only a few bristles showed distinguishable defects in *jbug^133^* homozygotes (Figure 2A). Phalloidin staining in *jbug^30^* pupae showed changes in F-actin staining consistent with the bent bristle phenotype (Figure 2B), suggesting that the bristle defects arise during pupation.

**Figure 2.**
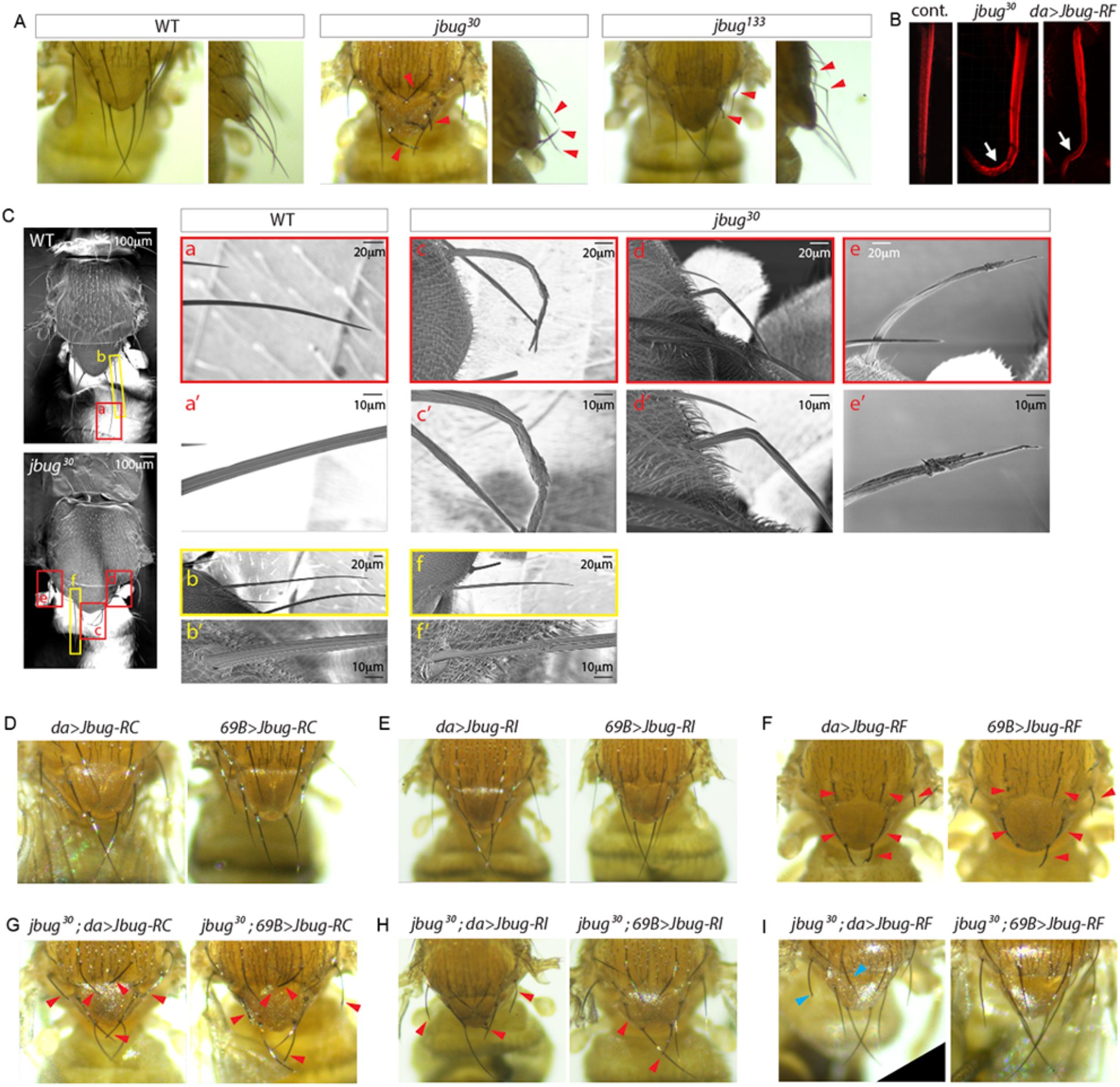
*jbug^30^* and *jbug^133^* show bristle defects on the adult thorax, which are rescued by expression of Jbug-RF. (A) Compared to wild type, *jbug^30^* and *jbug^133^* adult flies show distinct thoracic bristle defects, including short bristles with rough ends and bent bristles (arrowheads). *jbug^133^* flies have milder defects than *jbug^30^*. (B) Phalloidin staining (red) shows the bent bristle phenotype in *jbug^30^* and *da>Jbug-RF* pupae (arrows) compared to the straight wild-type bristle. (C) SEM images for bristles in wild-type and *jbug^30^* male flies. Bristles in the red and yellow boxes on the flies are shown in higher magnification in a-f. Compared to the nicely elongated and tapered wild-type bristle (a), bristles in *jbug^30^* mutants are short and distorted (c), bent (d), and rough-ended (e). Even when the bristle looks relatively normal, it is often shorter than the wild type and slightly twisted (compare b and f). (a’-f’) Higher magnification of a-f is shown. (D-F) Whereas overexpression of Jbug-RC (D) or Jbug-RI (E) using *69B-Gal4* or *da-Gal4* in an otherwise wild-type animal does not cause overt defects in thoracic bristles, overexpression of Jbug-RF causes short thick bristles (F; red arrowheads). (G-I) Although expression of Jbug-RC (G) or Jbug-RI (H) does not rescue the bristle defects in *jbug^30^* (red arrowheads), Jbug-RF expression almost completely rescues the bristle defects in *jbug^30^*, leaving only a couple of short bristles (I; cyan arrowheads).

To analyze the bristle defects in more detail, we performed scanning electron microscopy (SEM) in wild-type and *jbug^30^* homozygous mutant adults. Whereas wild-type animals showed nicely elongated and tapered bristles that suggested evenly aligned actin bundles (Figure 2C, a and a’), *jbug^30^* mutant flies had bristles that were short, curved (Figure 2C, c), bent (Figure 2C, d) and rough-ended (Figure 2C, e). Even when the bristles looked relatively normal, they were often shorter than wild type and mildly twisted (Figure 2C; compare b and b’ to f and f’).

Transheterozygotes of *jbug^30^* or *jbug^133^* over the *jbug^20^* null allele or the *59AB* deficiency line that deletes the entire *jbug* gene (Figure 1A) showed both semi-lethality and the same bristle defects as observed in homozygous *jbug^30^* and *jbug^133^* adults (Table 1). These data suggest that both phenotypes are due to loss of *jbug* function. Consistent with our data that *jbug^30^* is a more severe hypomorphic allele than *jbug^133^* (Figure 1, E and H), fewer transheterozygous *jbug^30^*/*jbug^20^* or *jbug^30^*/*59AB* flies survived to the adult stage compared to transheterozygous *jbug^133^*/*jbug^20^* or *jbug^133^*/*59AB* flies (Table 1). Taken together, these data suggest that Jbug-PF/M/N and/or PI are important for viability and actin bundle arrangement within bristles.

We also performed complementation tests using the deficiency line *Exel6079*, which deletes through the genomic region encoding the N-terminal exons of transcripts of *jbug-RL* and *jbug-RF/M/N* but leaves the coding region for *jbug-RC* and *jbug-RI* intact (Figure 1A). Interestingly, transheterozygous *jbug^30^*/*Exel6079* (and *jbug^133^*/*Exel6079*) flies showed neither semi-lethality nor bristle defects (Table 1). To test which isoforms are present in *jbug^30^*/*Exel6079* flies, we performed Western blot using extracts from *jbug^30^/Exel6079* transheterozygous 1^st^ instar larvae obtained from *jbug^30^* homozygous mothers and found that the truncated PI (∼90 Kd) band present in *jbug^30^* is no longer detected; only a full-length PI band is observed in the transheterozygotes. Also surprisingly, the large MW Jbug-PF/M/N bands that were absent/reduced in *jbug^30^* homozygotes were easily detected in *jbug^30^/Exel6079* larvae (Figure 1G). This finding was unexpected because Jbug-PF/M/N should be absent in transheterozygous *jbug^30^*/*Exel6079* flies; *jbug^30^* and *Exel6079* disrupt the 3’ and the 5’ ends of *jbug-RF/M/N*, respectively. Since *jbug^30^/Exel6079* larvae were obtained from *jbug^30^* homozygous mothers, Jbug-PI or Jbug-PF/M/N cannot be provided maternally. This result is challenging to reconcile, unless there is some form of transvection or transplicing that would generate full length transcripts coming from the 5’ regions of *jbug^30^* and 3’ regions of *Exel6079*. Such events could also explain the recovery of the full-length *jbug-RI* transcript and corresponding loss of the shortened form of Jbug-PI (Figure 1G).

### Expression of Jbug-RF rescues the bristle defects and semi-lethality of *jbug* hypomorphs

To begin to parse out the functions of the different Jbug isoforms, we created three new UAS lines encoding Jbug-PC (42 kDa protein), Jbug-PI (185.6 kDa) and Jbug-PF (>300 kDa protein). We also obtained a UAS construct for Jbug-PL (175.6 kDa protein) expression from the Mlodzik lab (Olguín *et al*. 2011). We first tested whether expression of any specific *jbug* splice form(s) could rescue the lethality of *jbug^20^* null mutants. Each UAS construct was expressed in *jbug^20^* mutant background using *daughterless-Gal4* (*da-Gal4; Wodarz et al., 1995*), which is expressed in all cells, or *69B-Gal4* (Rorth 1996), which is expressed in ectodermal cells. None of the UAS constructs rescued the lethality of *jbug^20^*; no larvae survived past the 1st instar larval stage as in *jbug^20^*. Co-overexpression of *jbug-RL* and *jbug-RI* together using da-Gal4 in the *jbug^20^* mutant background also did not rescue the lethality. These data suggest that multiple Jbug isoforms are required for viability.

We next tested whether expression of specific Jbug isoforms could rescue the bristle defects in *jbug^30^*, the severe hypomorphic allele. Consistent with the data that Jbug-PC is not affected in *jbug^30^* mutants, overexpression of *jbug-RC* with *da-Gal4* or with *69B-Gal4* in an otherwise wild-type background did not result in any overt defects in bristles (Figure 2D). Overexpression of Jbug-PI in an otherwise wild-type background also did not cause any defects in bristles (Figure 2E). On the other hand, overexpression of *jbug-RF* in otherwise wild-type animals resulted in shorter and thicker bristles (Figure 2F), suggesting that proper levels of Jbug-PF/M/N are critical for normal bristle development. Phalloidin staining in *fkh>Jbug-RF* pupae showed the bent bristle phenotype and irregular F-actin alignment, similar to observed in *jbug^30^* pupal bristles (Figure 2B). Importantly, whereas expression of *jbug-RC* (Figure 2G) or *jbug-RI* (Figure 2H) in *jbug^30^* mutants did not rescue the bristle phenotypes, expression of *jbug-RF* in *jbug^30^* mutants using *da-Gal4* or *69B-Gal4* nearly completely rescued the bristle phenotypes (Figure 2I). Moreover, *jbug-RF* expression in *jbug^30^* mutants also rescued semi-lethality. 86% (27/32) of pupae emerged as adult flies when *jbug-RF* was expressed in *jbug^30^* mutants, whereas only 26% (9/35) of *jbug^30^* homozygous pupae emerged as adult flies. These data suggest that the bristle defects and semi-lethality observed in *jbug^30^* are due to loss of the longest (>300 kDa) Jbug isoforms.

### Jbug hypomorphs show a delay in actin-based prehair formation and mild defects in wing hair orientation

Besides thoracic bristle defects, *jbug^30^* and *jbug^133^* homozygous adults also showed a mild swirling pattern of wing hairs (Supplemental Figure 3B and data not shown). Phalloidin staining of actin-rich prehairs at 32 hours after puparium formation (APF) revealed that hair formation is delayed in *jbug^30^* pupal wings (Supplemental Figure 3, D and E). A swirling pattern of wing hairs and delayed prehair formation are often observed in mutants for planar cell polarity (PCP) genes (Adler 2012). *jbug^30^* mutant eyes, however, did not have any defects in ommatidial rotation characteristic of loss of key PCP genes (Supplemental Figure 3, F and G), suggesting that Jbug is not involved in ommatidial rotation. Overall, these data suggest that proper levels for Jbug-PF/M/N and/or Jbug-PI are required for the timely maturation of the actin bundles that form the prehairs. In support of this, we find that the wing hair polarity defects are fully rescued in *jbug30/Exel6079* flies (Supplemental Figure 3C).

### Jbug proteins localize in a dynamic pattern in the embryonic epithelium

We next analyzed the subcellular localization of Jbug proteins during embryogenesis in wild type and *jbug* mutant alleles. During early development, Jbug was detected as small punctate structures in the cytoplasm of all cells (Figure 3, A and B). During stages 11-13, Jbug was detected at a high level both near the apical surface and at and near the adherens junctions (AJs) in all epidermal cells (Figure 3, C and M). AJ Jbug signals became weaker over time, and by stage 15, only mesh-like signals on the apical surface remained (Figure 3, D and M). In *jbug^20^* mutant embryos, AJ signals were absent, but signals on the apical surface were still detected (Figure 3, E and M). At stage 15 and later, apical surface signals were significantly decreased compared to wild-type levels (Figure 3, F and M). Since the *jbug^30^* and *jbug^133^* homozygotes that survived to adult stages were fertile, we analyzed Jbug signals in *jbug^30^* or *jbug^133^* embryos from homozygous mothers. Interestingly, in *jbug^30^* embryos from *jbug^30^* homozygous mothers, Jbug signals were both very weak and dispersed in epidermal cells throughout the stages and largely absent in AJs (Figure 3, G, H and M), suggesting that Jbug-PF/M/N and Jbug-PI might be major isoforms that show strong embryonic expression and AJ localization. Expression of Jbug-RF in the *jbug^30^* mutant background restored strong signals at AJs (Figure 3K) and on the apical surface (Figure 3L) at stage 13 and 15, respectively. Consistent with the minor changes in Jbug protein levels in *jbug^133^*, Jbug staining was mostly normal, albeit slightly reduced, in the epidermal cells in *jbug^133^* embryos from homozygous *jbug^133^* mothers (Figure 3, I, J and M).

**Figure 3.**
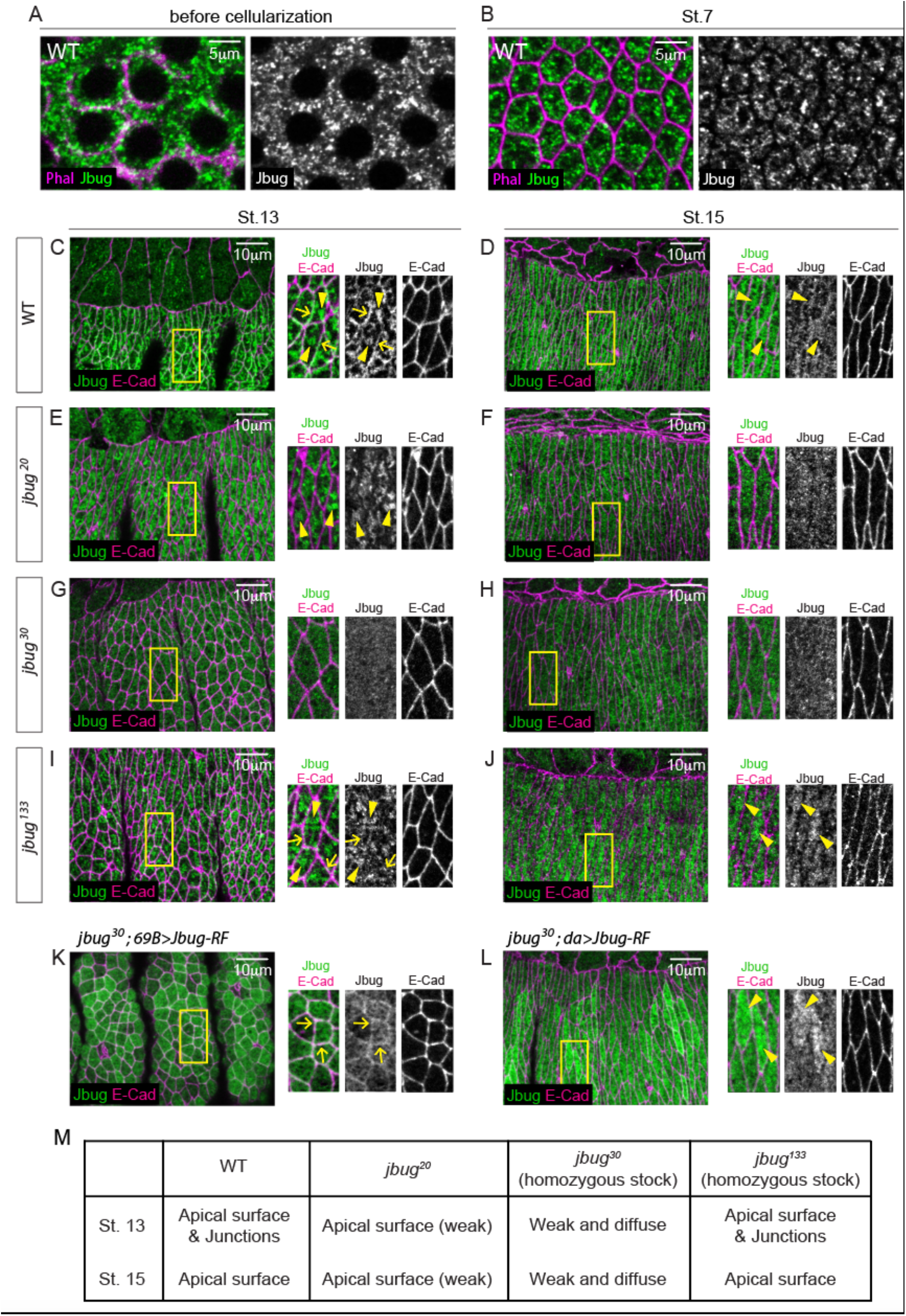
Analysis of Jbug protein localization in the embryonic epidermis in different *jbug* alleles. (A, B) Confocal images for early embryos stained for Jbug (green) and phalloidin (magenta). Jbug localizes as small apical punctate structures. (C-L) Confocal images for epidermis in stage 13 (C, E, G, I, K) and stage 15 (D, F, H, J, L) embryos stained for Jbug (green) and E-Cad (magenta). (C, D) In wild type, whereas high Jbug signals are observed both on the apical surface (arrowheads) and at adherens junctions (arrows) at stage 13 (C), Jbug mostly localizes more uniformly across the apical surface (arrowheads) at stage 15 (D). (E, F) In *jbug^20^* mutants, junctional Jbug signals are absent, and signals on the apical surface are weak (arrowheads in E). (G, H) In *jbug^30^* mutants, overall Jbug signals are very weak and diffused throughout the stages. (I, J) In *jbug^133^* embryos, signals at both the apical surface (arrowheads) and junctions (arrows) are present but are slightly weaker than wild type. (K, L) Jbug-RF overexpression in ectodermal cells driven by *69B-Gal4* in the *jbug^30^* mutant background (K) results in upregulation of Jbug signals in the entire apical domain with junctional enrichment at stage 13 (arrows). Overexpression of Jbug-RF in all cells by *da-Gal4* in the *jbug^30^* mutant background (L) causes upregulation of Jbug signals in the apical domain at stage 15 (arrowheads). (M) A summary of Jbug protein localization in the embryonic epidermis in wild type and *jbug* mutant alleles.

### Jbug is required for denticle formation in the epidermis

Jbug antibody staining in the ventral regions of stage 14 embryos revealed intense signals on the apical surface that presaged the actin-rich denticles that begin to form during stage 15 (Figure 4, B and C). This prompted us to ask if denticle formation was normal in *jbug* mutants. In wild-type embryos, six rows of epidermal cells in the anterior portion of each abdominal segment normally produce one or more denticles, forming a trapezoidal belt (Bejsovec 2013; Figure 4, C and D). Embryos homozygous for *jbug^20^* did not show any overt morphological defects, likely due to strong maternal contribution, but they exhibited defects in denticle formation (Figure 5B). Compared to wild type (Figure 5A), many denticles in *jbug^20^* embryos were small and premature (Figure 5B). Also, the overall number of denticles were fewer in *jbug^20^* embryos compared to wild type. We counted the number of denticles in each cell from a grid of 24 cells (Figure 5K) for at least five individuals of each genotype. *jbug^20^* embryos had fewer denticles per cell than wild-type embryos in several rows of cells (C1, C3, C4 and C6; Figure 5, L and M; Supplemental Figure 4, A and B). Denticles in *jbug^30^* embryos from a homozygous stock were relatively normal in their size and morphology, but these embryos had significantly fewer denticles compared to wild type (Fgure 5C). Irrespective of which row of cells (C1-C6) was examined, *jbug^30^* embryos had, on average, fewer denticles per cell than wild-type embryos (Figure 5, L and M; Supplemental Figure 4, A and B). These data suggest important roles of Jbug in actin bundle arrangement for denticle formation.

**Figure 4.**
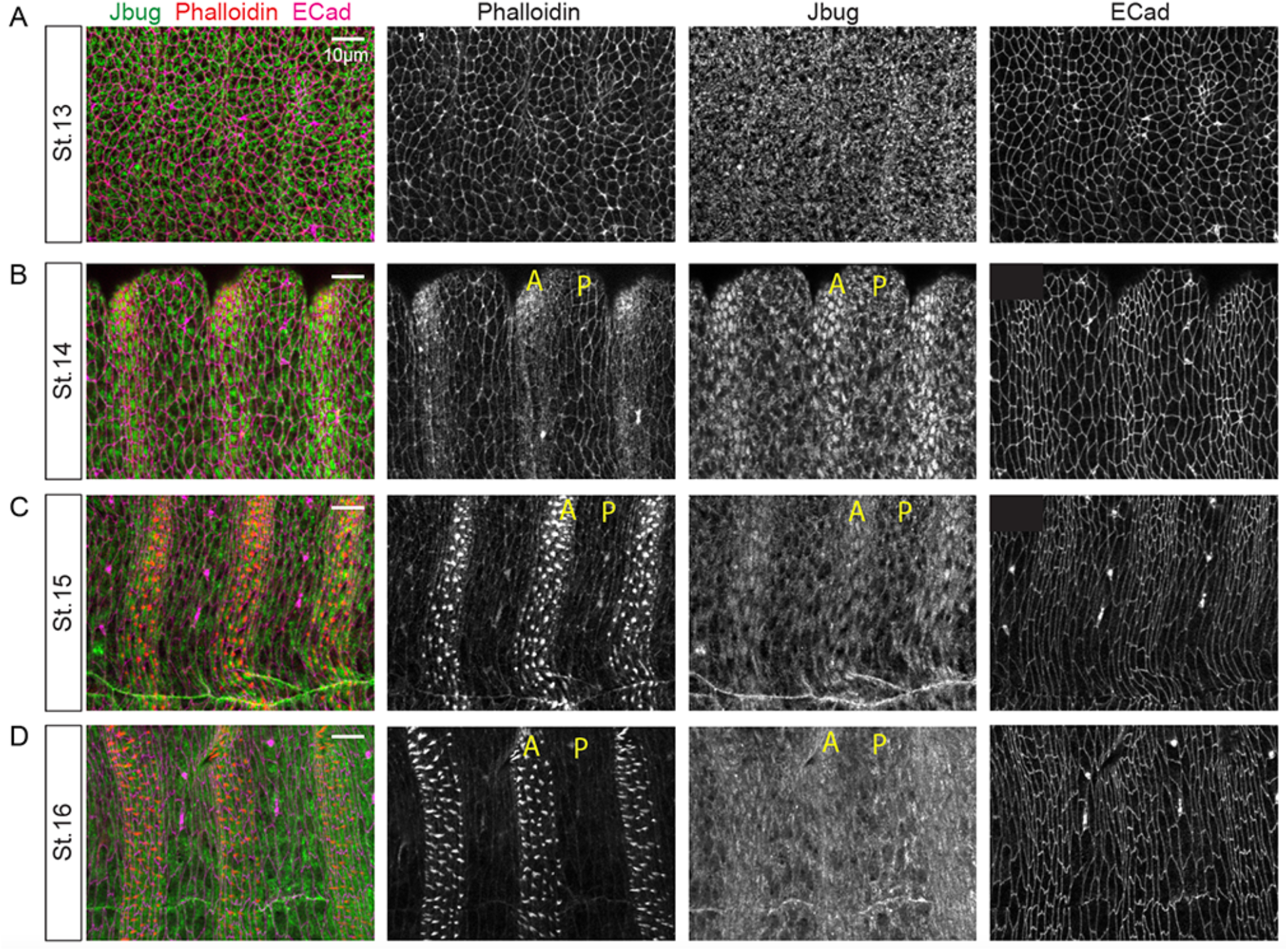
Jbug protein is upregulated in the anterior part of each segment just prior to the formation of actin-rich denticles in the same region. (A-D) Epidermis in stages 13 (A), 14 (B), 15 (C) and 16 (D) embryos stained for Jbug (green), phalloidin (red), and E-Cad (magenta). (A) At stage 13, Jbug is mostly uniform across epidermis. (B) At stage 14, actin begins to be upregulated in the anterior side of each segment. Jbug also shows accumulation in the apical domain of epidermal cells in the anterior side of the segment. (C-D) At stages 15 (C) and 16 (D), actin-rich denticles form in the anterior side of each segment. Jbug still shows higher levels in those regions. A and P represent the anterior and the posterior side, respectively.

**Figure 5.**
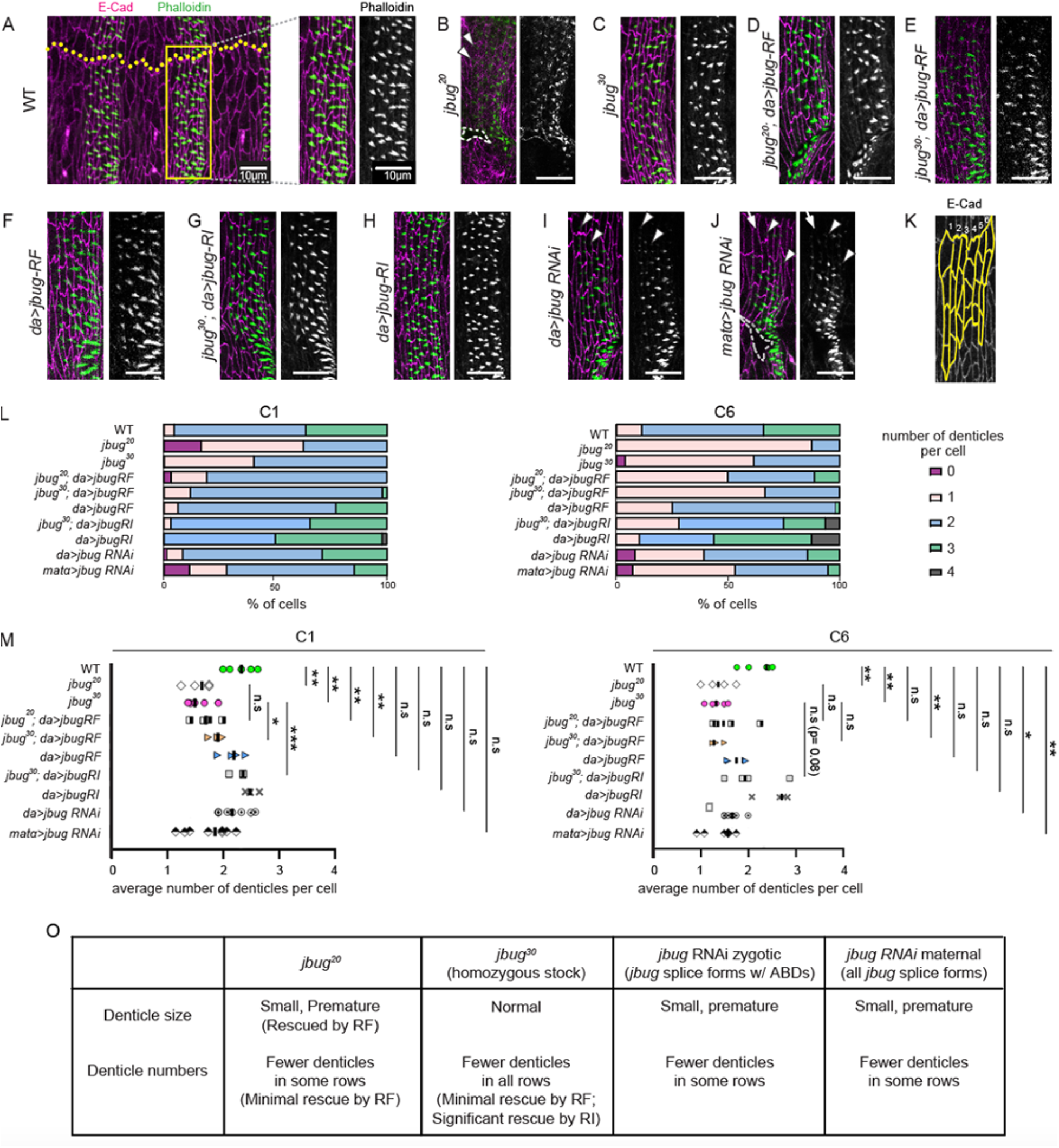
Jbug regulates the formation of denticle precursors in the ventral epidermis of the embryo. (A-G) Stage 16 embryos stained for E-Cad (magenta; cell boundaries) and phalloidin (green; denticles). Ventral epidermis in wild type (A), *jbug^20^* (B), *jbug^30^* (C), *jbug^20^*; *da-Gal4>Jbug-RF* (D), *jbug^30^*; *da-Gal4>Jbug-RF* (E), *da-Gal4>Jbug-RF* (F), *jbug^30^*; *da-Gal4>Jbug-RI* (G), *da-Gal4>Jbug-RI* (H), *da-Gal4>jbug RNAi* (*GD13033*) (I) and *matα-Gal4>jbug RNAi* (*JF01166*) (J) is shown. (A) Epidermal cells at the denticle belt in the abdominal segment 5 (yellow box) are shown at higher magnification. Dotted yellow line, ventral midline. (B-J) Epidermal cells of the same region as in A are shown. For all the embryos, anterior is to the left. (B, J) *jbug^20^* mutants (B) and *matα-Gal4>jbug RNAi* embryos (J) show mild defects in the embryo morphology, such as wavy epidermis (areas with dotted lines; cells are out of focus). Occasionally, cells either fail to form denticles (arrow in J) or form very short denticles (arrowheads in B, I and J). (K) A cartoon for quantification of the denticle numbers. Four cells that are closest to the ventral midline were chosen in each of the six columns of cells in denticle belts in abdominal segments 4, 5 and 6 (column 1, most anterior). (L) Percentage of cells that have different numbers of denticles in C1 and C6. See Supplemental Figure 4 for quantification for C2-C5. (M) Quantification of the average number of denticles per cell in C1 and C6. Student’s t-test with Welch’s correction is used for statistics calculation. (O) A summary of denticle defects in *jbug* mutant alleles and RNAi knockdown and rescue results with Jbug-RF and -RI.

We next tested whether expression of *jbug-RF* or *jbug-RI* rescues the denticle defects in *jbug^20^* or *jbug^30^* embryos. Expression of *jbug-RF* using the *da-Gal4* driver in the otherwise wild-type condition did not affect the number of denticles nor the size (Figure 5, F, L and M). Expression of *jbug-RF* using the *da-Gal4* driver in *jbug^20^* mutants largely restored denticle size (compare Figure 5B and 5D), suggesting a role of the Jbug isoforms with actin-binding domains in formation of mature denticles. However, Jbug-RF expression in *jbug^20^* minimally rescued the decreased denticle number phenotype; it slightly increased the number of denticles in one row (C6), but not significantly in other rows (C1-C5) (Figure 5, D, L and M; Supplemental Figure 4, A and B). Similarly, expression of *jbug-RF* using the *da-Gal4* driver in *jbug^30^* mutants slightly increased the number of denticles in one row (C1), but not in other rows (C2-C6) (Figure 5, E, L and M; Supplemental Figure 4, A and B). On the other hand, overexpression of *jbug-RI* using the *da-Gal4* driver in the otherwise wild-type condition significantly increased the number of denticles in several rows (C2-C4; Supplemental Figure 4, A and B). In all rows, Jbug-RI-overexpressing embryos have more cells with more than three denticles per cell than wild type (Figure 5L; Supplemental Figure 4, A and B). Moreover, expression of *jbug-RI* using *da-Gal4* in *jbug^30^* mutants significantly rescued the decreased denticle number phenotype in several rows (Figure 5, G, L and M; Supplemental Figure 4, A and B). These data suggest a critical role of Jbug-PI in formation of the actin-rich denticles in the embryo.

We further tested whether the maternal pool of *jbug* is required for denticle formation. Knocking down *jbug-RL* and *jbug-RF/M/N* by RNAi (*GD13033*; Figure 1A) using *da-Gal4* also showed similar, albeit weaker, defects to those in *jbug^20^* embryos: a decrease in the size and numbers of denticles per cell in several rows (Figure 5, I, L and M; Supplemental Figure 4, A and B), suggesting roles for the Jbug isoforms with actin-binding domains in denticle formation. Maternal knockdown of *jbug* using the *matα-Gal4* driver (Häcker and Perrimon 1998) and two strong RNAi lines (*GD8664* and *GD13033*; each targets a subset of *jbug* splice forms; Figure 1A) resulted in very few embryos, suggesting an essential role for Jbug in oogenesis. Maternal knockdown of *jbug* using the weaker RNAi line (*JF01166* that targets all splice forms; Figure 1A) resulted in mostly normal overall embryonic morphology, except for slightly wavy epidermis (Figure 5J), and these embryos showed defects in denticle formation. As observed in *jbug^20^* and *jbug^30^* mutants or *da-Gal4>jbug RNAi GD13033* embryos, some cells failed to form denticles; others formed short, premature denticles with weak phalloidin signals (Figure 5, J, L and M). Quantification showed a decrease in the average number of denticles per cell in most columns (except for C2) of each denticle belt with *matα*>*jbug RNAi* (Figure 5, L and M; Supplemental Figure 4, A and B). Taken together, our data suggest that both maternally and zygotically provided Jbug proteins are required for denticle formation.

### Jbug is upregulated in embryonic epithelial tubes

Consistent with mRNA expression (BDGP; https://insitu.fruitfly.org/cgi-bin/ex/insitu.pl; Fly-FISH; http://fly-fish.ccbr.utoronto.ca; Supplemental Figure 1), antibody staining in wild-type embryos revealed accumulation of Jbug protein in the trachea and the salivary gland (Figure 6, A and E). Interestingly, Jbug was abundant in both the cytoplasm and in a distinct apical domain of both tracheal and salivary gland cells, which appeared to be just under the apical surface (Figure 6, A and E), as opposed to following the AJ pattern of E-Cad. This apical staining in the salivary gland was prominent up to stage 14; at later stages, Jbug signals were observed mostly in the cytoplasm and detected in the apical domain only as a thin layer along the lumen (compare Figure 6E to Figure 7A).

**Figure 6.**
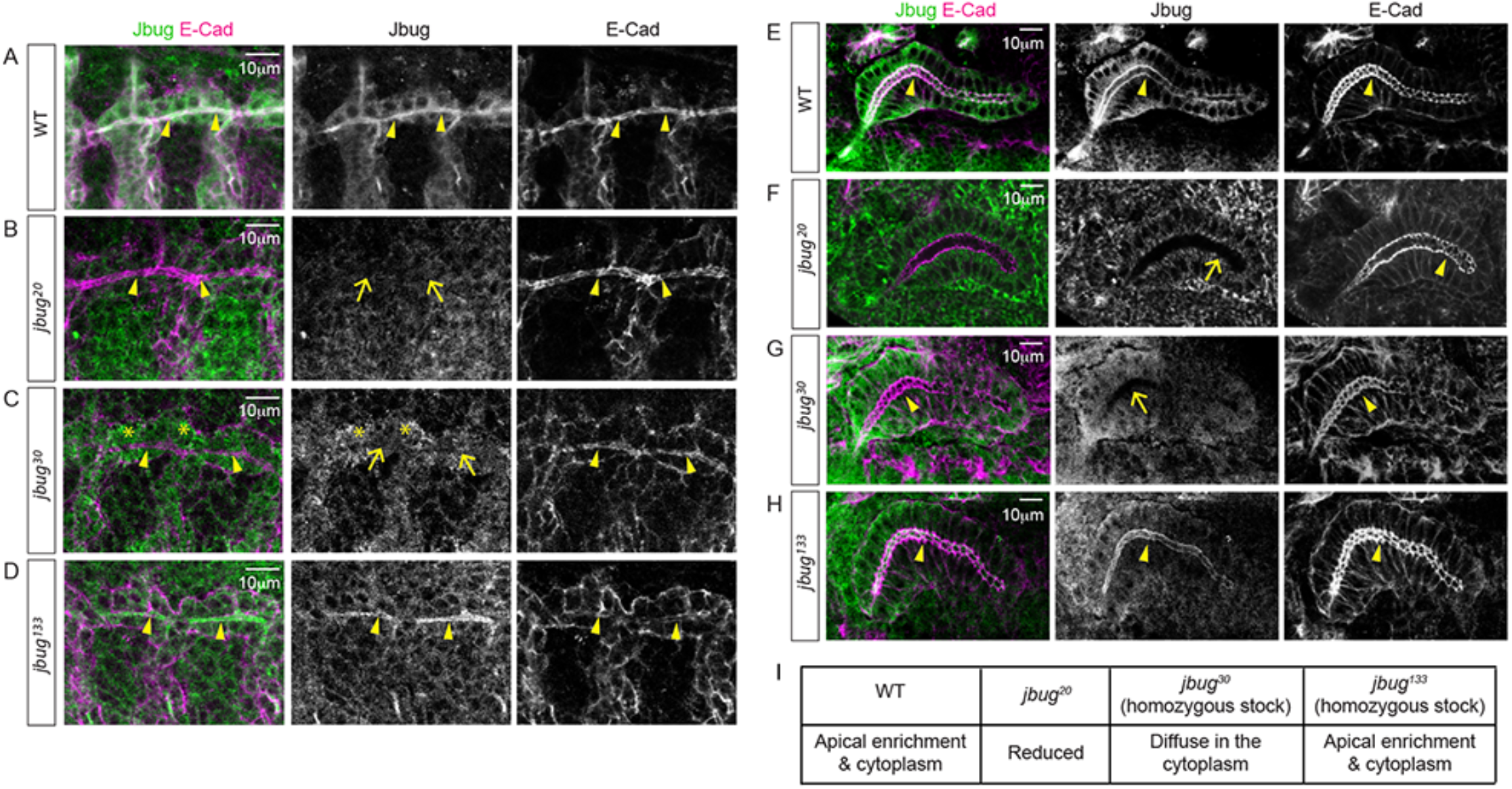
Jbug protein is apically enriched in the embryonic trachea and the salivary gland, and this enrichment is absent in *jbug^20^* and *jbug^30^* mutants. (A-D) Confocal images of stage 14 embryonic trachea stained for Jbug (green) and E-Cad (magenta). (A) In wild type, weak Jbug signals are observed in all tracheal cells, with enrichment in the apical domain (arrowheads). (B) In *jbug^20^* embryos, Jbug signals in the trachea are mostly absent (arrows). (C) In *jbug^30^* mutant embryos, Jbug signals are diffused in the cytoplasm in tracheal cells (asterisks), with no apical enrichment (arrows; compare to E-Cad signals (arrowheads) in the apical domain). (D) In *jbug^133^*, apical enrichment of Jbug is still present (arrowheads). (E-H) Stage 14 salivary glands stained for Jbug (green) and E-Cad (magenta). (E) Similar to tracheal signals, Jbug is enriched in the apical domain of wild-type salivary gland cells (arrowheads) with weak signals in the cytoplasm. (F) In *jbug^20^*, overall Jbug signals are significantly decreased in the salivary gland and apical enrichment is mostly gone (arrow; compare to E-Cad signals (arrowhead) in the apical domain). (G) In *jbug^30^*, Jbug is not apically enriched in the salivary gland (arrow; compare to E-Cad signals (arrowhead) in the apical domain) but diffused in the cytoplasm. (H) In *jbug^133^*, apical enrichment of Jbug is still present (arrowheads). (I) A summary of Jbug protein localization in the trachea and the salivary gland in wild type and *jbug* mutant alleles.

**Figure 7.**
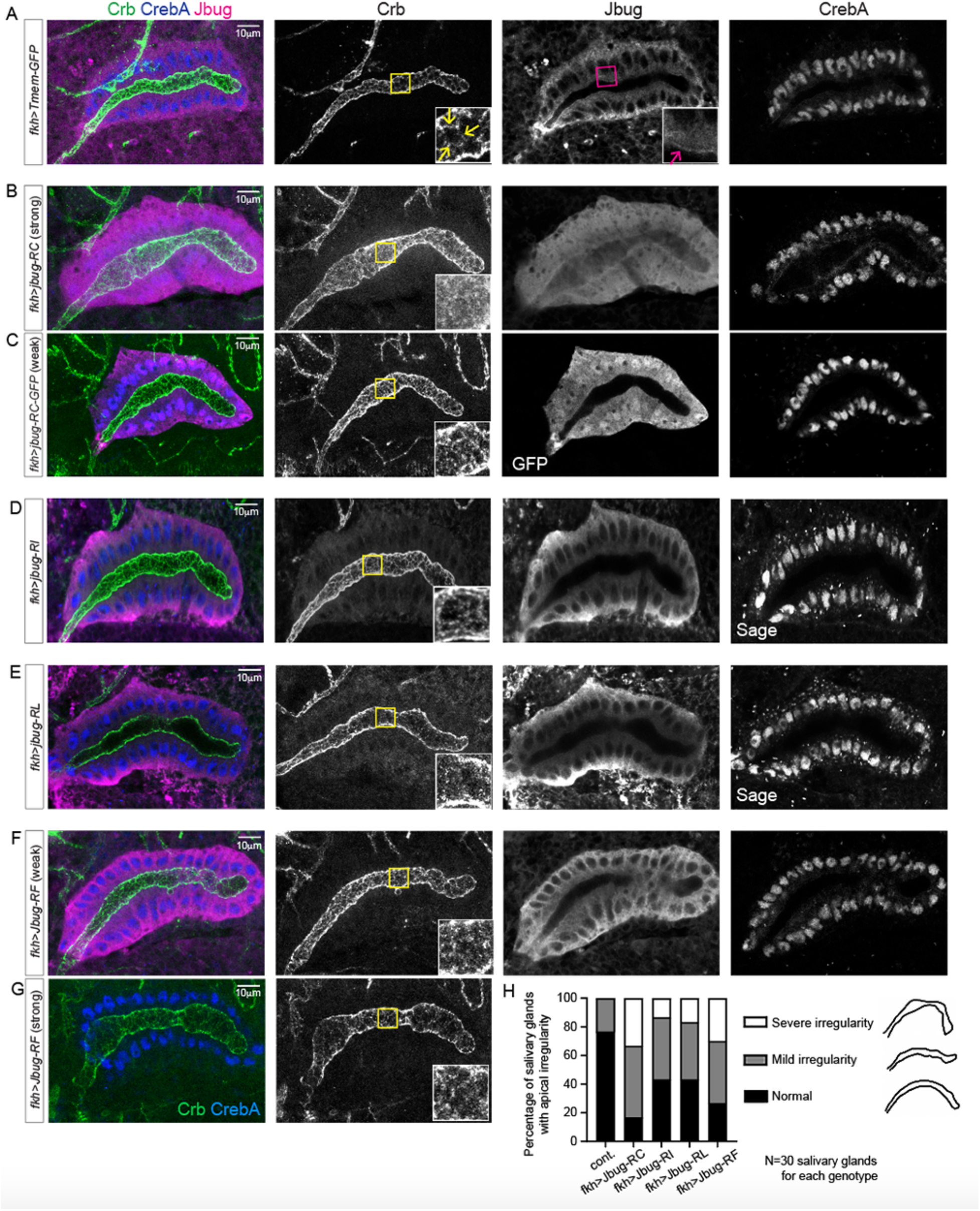
Jbug overexpression in the salivary gland causes mislocalization of Crb along the apical surface and an expanded and irregular apical domain. (A-D) stage 16 salivary glands stained for Crb (green), Jbug (magenta), GFP (magenta in C) and salivary gland nuclear markers CrebA (A-C and F) and Sage (D and E). Salivary glands overexpressing Tmem-GFP (control, A), Jbug-RC (B), Jbug-RC-GFP (C), Jbug-RI (D), Jbug-RL (E) and Jbug-RF (F, G) are shown. Note that Jbug mainly localizes to the cytoplasm and only shows weak apical signals in stage 16 salivary glands (red arrow in A; compare to Figure 4E). Jbug-RC (B) and Jbug-RC-GFP (C) localize both to salivary gland nuclei and the cytoplasm when overexpressed. Jbug overexpression causes an expanded and irregular salivary gland lumen (B-G). Irregular salivary gland lumen is worsened when stronger UAS lines are used (compare B and C; also compare F and G). Compared to subapical Crb signals along AJs in control salivary gland cells (inset in A; yellow arrows), Crb signals are dispersed along the entire apical domain in Jbug-overexpressing salivary glands (insets in B-G). (H) Quantification of the irregular apical membrane morphology in salivary glands overexpressing different Jbug isoforms. Strong UAS lines shown in B and G were used to overexpress Jbug-RC and Jbug-RF. For mild irregularity, salivary glands with the uneven apical domain were counted. For severe irregularity, salivary glands with bulging and constriction phenotypes were counted.

Jbug signals in tubular epithelial organs were significantly decreased in *jbug^20^* embryos, even with regards to surrounding tissues (Figure 6, B and F), suggesting that zygotic *jbug* expression is required for upregulation of Jbug in these tissues. In *jbug^20^*, the apical staining in the trachea was absent and in the salivary gland was more punctate. In *jbug^30^* embryos (from *jbug^30^* parents), apical staining of Jbug in the trachea and salivary glands was also absent; instead, uniform Jbug signals were detected in the cytoplasm (Figure 6, C and G). Consistent with slight changes in Jbug protein levels in *jbug^133^*, Jbug signals were mostly normal, albeit slightly reduced, in the trachea and salivary glands in *jbug^133^* embryos from *jbug^133^* parents (Figure 6, D and H). These data suggest that it is either the Jbug-PF/M/N and/or Jbug-PI that are enriched in the apical domain in tubular epithelial organs in the embryo.

Despite the absence or mislocalization of Jbug in the apical domain of trachea and salivary glands, both organs developed normally without overt morphological defects in *jbug^20^* and *jbug^30^* embryos (Figure 6, B, C, F and G). Crumbs (Crb), the apical determinant protein, E-Cadherin (E-Cad), a key adherens junction protein, Discs large (Dlg), a septate junction component, and α-Spectrin (α-Spec), the lateral membrane-associated protein, also localized properly in salivary glands in *jbug^20^* or *jbug^30^* embryos (Supplemental Figure 5, A-I), suggesting that the overall epithelial polarity is intact in salivary glands in these mutants. Also, phalloidin signals in salivary glands in *jbug^20^* and *jbug^30^* embryos were comparable to wild type (Supplemental Figure 5, D-F), suggesting that F-actin is largely intact in salivary glands in *jbug* mutants.

To further test where each of the jbug isoforms localizes and to test the effect of Jbug overexpression in tubular epithelial tissues, we overexpressed the different Jbug isoforms (Jbug-PC, PI, PL and PF) in the salivary gland using the salivary gland-specific *fork head-Gal4* driver (*fkh-Gal4*; Henderson and Andrew 2000). All isoforms primarily localized in the cytoplasm when overexpressed (Figure 7, D-G). Interestingly, however, both untagged and C-terminal GFP-tagged Jbug-PC were detected both in the cytoplasm and in nuclei (Figure 7, B and C). These data suggest that this smallest isoform – which is primarily expressed in pupae (Figure 1E) – might function in the nucleus as well. Similar nuclear localization has been observed with human Filamin A. Although full-length filamin A is predominantly cytoplasmic, a C-terminal fragment of Filamin A colocalizes with the androgen receptor to the nucleus to act as a co-transcription factor (Loy *et al*. 2003). Overexpression of each isoform resulted in an expanded and irregular luminal domain (Figure 7, B-G). Interestingly, Crb was mislocalized and dispersed along the entire apical surface in Jbug-overexpressing salivary glands (Figure 7, B-G) rather than being enriched in a sub-apical domain just apical to the AJs as in control salivary glands (Fig 7A, inset in Crb column; Wodarz *et al*. 1995; Tepass 1996; Chung and Andrew 2014). These data indicate that overexpression of multiple Jbug isoforms can disrupt the sub-apical domain enrichment of Crb. Phalloidin and other junctional and polarity markers, including E-Cad, Nrx-IV (a septate junction marker) as well as α-Spec, localized properly in Jbug-overexpressing salivary glands (Supplemental Figure 6), suggesting that the overall epithelial polarity is not disrupted by Jbug overexpression. Taken together, these data suggest that appropriate levels of Jbug isoforms are important for proper Crb localization and apical organization of epithelial cells during embryogenesis.

### Jbug is required for maintaining Crb levels in embryonic epithelial cells

Maternal knockdown (*matα*>*JF01166*) or zygotic loss of all *jbug* splice forms (*jbug^20^*), or specific maternal and zygotic loss of the *jbug-RF/M/N* and *jbug-RI* (*jbug^30^*) did not show overt morphological defects in the embryo, except for mild wavy epidermis (Figures 5, B and J) and denticle formation defects (Figure 5). We therefore knocked down both maternal and zygotic pools of all *jbug* splice forms using RNAi. Compared to normal morphology in control embryos (Figure 8, A and E) or in embryos knocked down for *jbug* zygotically (using *da-Gal4*; Figure 8, B and F) or maternally (using *matα*-Gal4; Figure 8, C and G), knockdown of *jbug* both maternally and zygotically using *matα-Gal4* and *da-Gal4* (*jbug* MZ knockdown) resulted in a severely disrupted morphology in all embryos produced (Figure 8, D and H). The embryos were often so disrupted that their stages could not be ascertained. Whereas strong Jbug signals were still detected in epithelial cells in embryos knocked down for *jbug* maternally or zygotically (Figure 8, F’’ and G’’; compare to control in Figure 8E’’), Jbug signals were absent in the apical surface in many epidermal cells in *jbug* MZ knockdown embryos (Figure 8H’’). Importantly, Crb signals were also absent in those cells (Figure 8H’’’). These data suggest a critical role for Jbug in regulating Crb levels and localization in epithelial cells.

**Figure 8.**
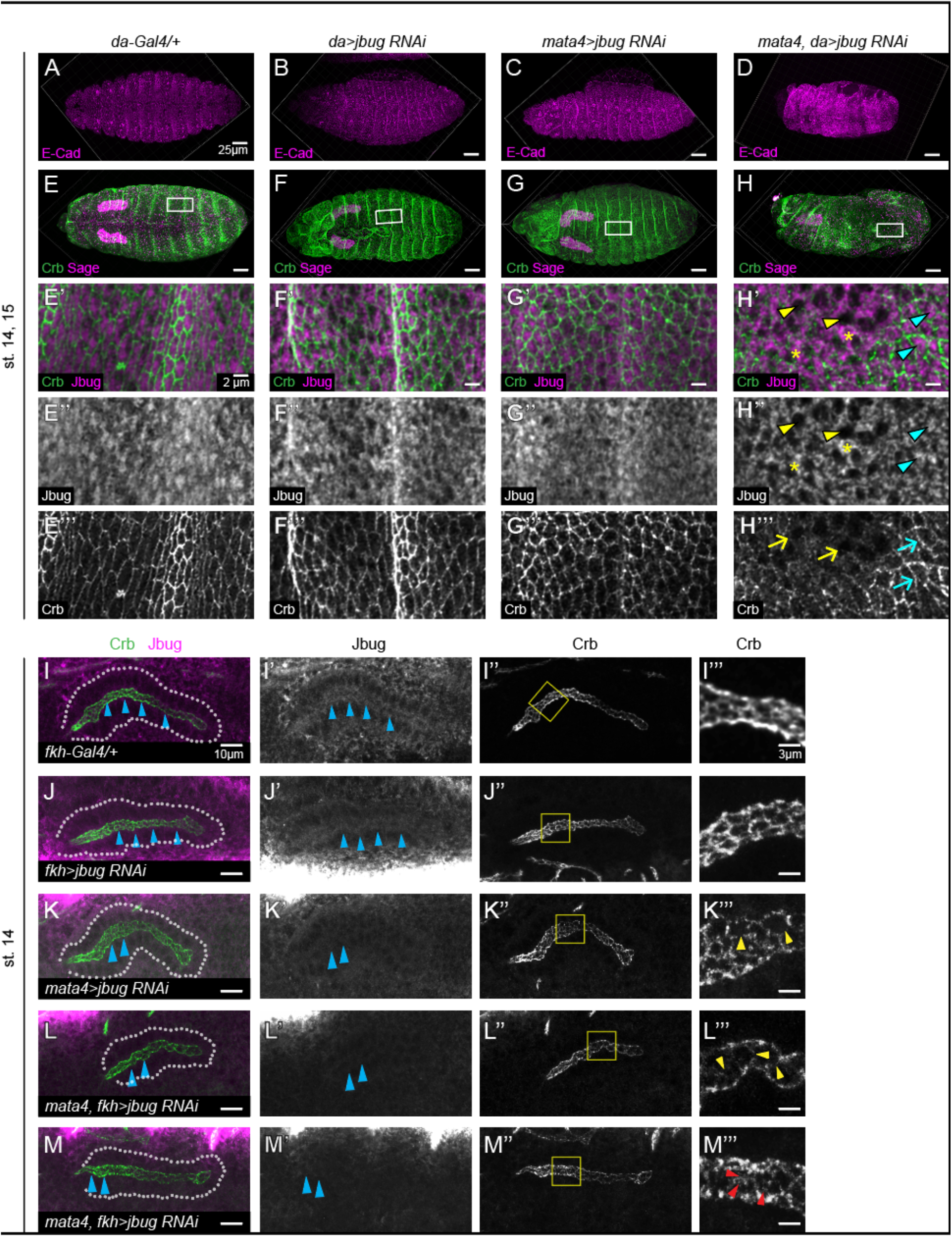
Jbug is required for proper Crb levels in epithelial cells during embryogenesis. (A-H) Confocal images of stage 14/15 embryos of control (*da-Gal4/+*; A and E), zygotic (*da>jbug RNAi*; B and F), maternal (*matα>jbug RNAi*; C and G), and both maternal and zygotic knockdown of *jbug* (*matα, da>jbug RNAi*; D and H). Anterior is to the left. Ventral (A, D, E), dorsal (F) and lateral (B, C, G and H) views are shown. (A-D) Embryos stained for E-Cad (magenta). (E-H) Embryos stained for Crb (green) and Sage (a SG marker; magenta). (E’-H’’’) Higher magnification of the boxed regions in E-H. Crb (green) and Jbug (magenta) are shown. Compared to strong apical surface Jbug signals in epidermal cells in control (E’’) and embryos knocked down for *jbug* only zygotically (F’’) or maternally (G’’), Jbug signals are absent in the apical surface in many epidermal cells in *jbug* M/Z knockdown embryos (yellow arrowheads in H’ and H’’). In those cells, Crb is also absent (yellow arrows in H’’’). Compare to neighboring cells with significant apical surface Jbug signals (cyan arrowheads in H’ and H’’) and relatively normal Crb localization (cyan arrows in H’’’). Asterisks, residual or nonspecific Jbug signals. (I-M) Confocal images of stage 14 salivary glands stained for Crb (green) and Jbug (magenta). White dotted lines, salivary gland boundaries. Control (*fkh-Gal4/+*; I), zygotic (*fkh>jbug RNAi*; J), maternal (*matα>jbug RNAi*; K), and both maternal and zygotic knockdown of *jbug* (*matα, fkh>jbug RNAi*; two representative images are shown in L and M). (I’-M’’) Jbug (I’-M’) and Crb (I’’-M’’) only. Jbug signals are reduced in *jbug* knockdown (cyan arrowheads in J’-M’), with a more significant reduction in maternal (K’) and maternal/zygotic knockdown (K’-M’). (I’’’-M’’’) Higher magnification of the boxed regions in I’’-M’’. Maternal and maternal/zygotic knockdown of *jbug* results in gaps (yellow arrowheads in K’’’ and M’’’) or mislocalized Crb signals to the apical surface (red arrowheads in L’’’).

We also knocked down *jbug* in the salivary gland in embryos also maternally knocked down for *jbug,* using *matα-Gal4* and *fkh-Gal4*. No overt morphological defects were observed in salivary glands in these embryos at stage 11, when the salivary gland begins to invaginate to form a tube (Supplemental Figure 7). However, compared to continuous Crb signals in control or in salivary glands knocked down for *jbug* maternally or zygotically only, huge gaps in Crb signals were detected in salivary glands in *jbug* MZ knockdown embryos (Supplemental Figure 7, A’’’’-D’’’’). At later stages, knockdown of *jbug* resulted in a slightly irregular salivary gland lumen (Figure 8, I-M). Importantly, compared to the continuous sub-apical localization of Crb in control or SG cells with zygotic knockdown for *jbug*, the salivary glands knocked down for *jbug* maternally showed discontinuous Crb signals (Figure 8I’’’). Salivary glands knocked down for *jbug* both maternally and zygotically showed reducted Crb signals with wider gaps (Figure 8L’’’) or mislocalization of Crb signals to the apical surface (Figure 8M’’’). Overall, these data reveal that Jbug is required for proper levels and distribution of Crb in epithelia.

## Discussion

Filamins play important roles in actin reorganization in many developmental processes and disease contexts. In this study we show that different isoforms of Jbug, a *Drosophila* filamin-type protein, are differentially expressed and localized during development. Using new genetic tools that we generated, we reveal essential roles for different Jbug isoforms in viability, epithelial morphogenesis and formation of actin-rich structures.

### Roles of different Jbug isoforms in various developmental stages in *Drosophila*

Our developmental western analysis (Figure 1), along with developmental RNA-Seq data by Flybase (www.flybase.org; FB2021_02), reveals differential expression of different Jbug isoforms throughout development. Although transcripts of isoforms containing actin-binding domains (*jbug-RF/M/N* and *jbug-RL*) are not expressed until several hours of embryogenesis, proteins for these isoforms are detected at early hours of embryogenesis, suggesting Jbug proteins are provided maternally. Our studies reveal roles of different Jbug isoforms at various developmental processes, including oogenesis, epithelial morphogenesis, and viability at larval and pupal stages.

Severe defects in egg laying upon maternal knockdown of *jbug* using two strong RNAi lines that target a different subset of isoforms (*GD13033* and *GD8664*) suggest critical roles for Jbug isoforms in oogenesis. *GD13033* targets Jbug-PF/M/N and Jbug-PL, suggesting roles for the actin-binding domain-containing isoforms in oogenesis. Homozygous *jbug^30^* flies can be maintained as a homozygous stock depite the apparent absence of Jbug-PF/M/N and truncated Jbug-PI, suggesting that these Jbug isoforms do not play an essential role in oogenesis. Since Jbug-PC is only made in pupae, <u>the main isoform for oogenesis is likely to be Jbug-PL</u> (Figure 9).

**Figure 9.**
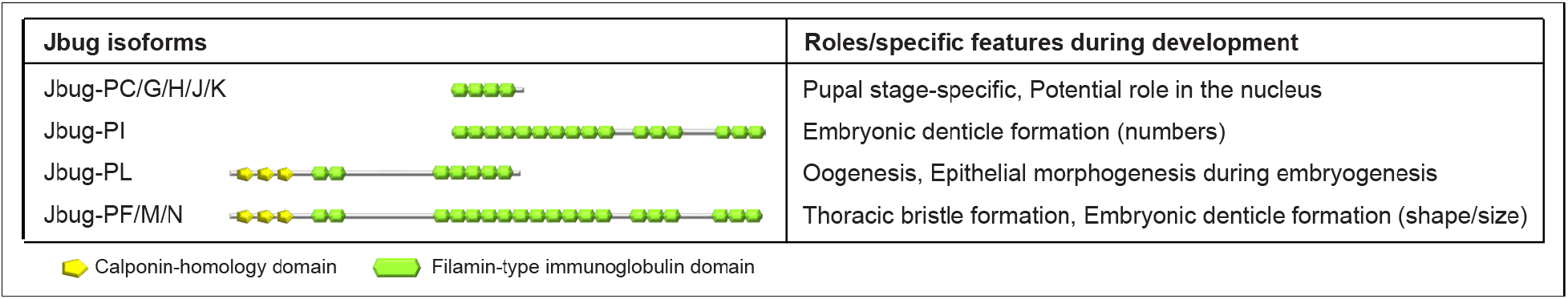
Isoform-specific roles and features of Jbug.

Our study also reveals a critical role of Jbug in epithelial morphogenesis during embryogenesis. Embryos knocked down for *jbug* both maternally and zygotically show severe morphological defects with disrupted Crb localization (Figure 8). The precise mechanism underlying Crb regulation by Jbug remains to be discovered. As filamin binds to many interacting proteins via IG-FLMN repeats, it will be interesting to test whether Jbug regulates Crb by directly interacting with it. It is also possible that Crb reduction is an indirect effect of disrupted actin cytoskeleton by loss of Jbug. Dynamic localization of Jbug isoforms to the apical surface during embryogenesis (Figures 3 and 4) suggests potential roles for different Jbug isoforms in remodeling the actin cytoskeleton and contributing to the mechanical stability of the plasma membrane and the cell cortex. Interestingly, homozygous *jbug^30^* embryos from homozygous mothers, where only Jbug-PC and Jbug-PL are intact, show no overt defects during embryogenesis except for defects in denticle formation at later stages (Figure 5). As Jbug-PC is not expressed during embryogenesis (Figure 1), these data indicate that <u>Jbug-PL is largely sufficient for epithelial morphogenesis during embryogenesis</u> (Figure 9).

Jbug is also required for viability. The *jbug^20^* null allele lacking all Jbug isoforms is 1^st^ instar larval lethal. 1^st^ instar larvae of homozygous *jbug^20^* or transheterozygous animals of *jbug^20^* and a deficiency line appear normal in their overall size, morphology and motility, but they do not molt into the 2^nd^ instar. Based on the role of Jbug in regulating Crb localization during embryogenesis (Figures 7 and 8), the lethality of *jbug^20^* homozygous larvae might be linked to disruption of Crb localization and/or epithelial organization during the larval stage. As overexpressing a single isoform or Jbug-RL and Jbug-RI together failed to rescue the lethality of *jbug^20^*, multiple Jbug isoforms may be required for viability during early stages of development.

Our *jbug* hypomorphic alleles reveal roles for Jbug-PF/M/N and/or Jbug-PI in organismal viability late in development (Table 1). Most flies homozygous for *jbug^30^* or *jbug^133^*, or transheterozygous flies for these *jbug* alleles and a deficiency completely removing *jbug* die during late pupation. Jbug-PF expression in the *jbug^30^* mutant background rescued this semi-lethality, suggesting that the longest splice forms (PF/M/N) are required at these late stages. Interestingly, anatomy RNA-Seq by Flybase (www.flybase.org; FB2021_02) shows increased *jbug* expression in the fat body at pupal stage P8 (2 days old; moderate levels of expression) compared to the white prepupae stage (very low expression), opening up the possibility for a role for Jbug in the fat body. The fat body of *Drosophila* is equivalent to vertebrate adipose tissue and liver in its storage and major metabolic functions, including immune response and sensing nutritional conditions. It remains to be revealed whether the lethality caused by the loss (or significant reduction) of Jbug-PF/M/N is related to defects in morphogenesis, metabolic functions, or both.

### Jbug plays a role in formation of actin-rich protrusions

Our results identify roles for specific Jbug isoforms in formation of two prominent actin-rich protrusions in *Drosophila*, bristles on the thorax and denticles in the ventral epidermis of the embryo (Figures 2 and 5). Jbug may regulate assembly of actin bundles by interacting with actin-regulating proteins, such as capping protein and the Arp2/3 complex components. Mutants for capping protein or the Arp2/3 complex components show bristle defects reminiscent of the *jbug* mutant phenotypes (Hudson and Cooley 2002; Frank *et al*. 2006). Several studies in other systems also suggest that the Arp2/3 complex synergizes with filamins. In filamin-deficient melanoma cells, the Arp2/3 complex is still abundant but unable to stabilize the plasma membranes or support locomotion (Cunningham *et al*. 1992). Also, branched actin filaments made with Arp2/3 do not have the closing loop structures necessary to form three-dimensional structures (Mullins *et al*. 1998; Blanchoin *et al*. 2000), unless the filaments are assembled in the presence of filamin (Niederman *et al*. 1983; Wolosewick and Condeelis 1986). Other good candidates that might interact with Jbug to organize the actin cytoskeleton include the Rho family small GTPases and their regulators. The Rho family small GTPases, including Rho, Rac, and Cdc42 are known to physically interact with Filamin (Ohta *et al*. 1999; Jeon *et al*. 2008; Del Valle-Pérez *et al*. 2010), as are regulators of Rho GTPases, including guanine nucleotide exchange factors (Bellanger *et al*. 2000; Pi *et al*. 2002), GTPase activating proteins (Ohta *et al*. 2006; Mammoto *et al*. 2007; Nakamura *et al*. 2009) and Rho kinase (Ueda *et al*. 2003). It will be interesting to test whether Jbug works with any of these proteins to regulate bristle and/or denticle formation.

Defects in actin-rich protrusions in *jbug^30^* are observed in the presence of Jbug-PL, which contains actin-binding domains and short IG-FLMN repeats, suggesting that this isoform is not sufficient to form normal bristles and denticles in *Drosophila*. Indeed, bristle defects are rescued by expression of Jbug-PF, one of the longest isoforms that contain long IG-FLMN repeats following the actin-binding domains (Figure 2). <u>These data suggest an important role for protein interactions through the long IG-FLMN repeats in forming actin-rich protrusions</u>. Our interpretation is consistent with a role of filamins in acting as a molecular scaffold. More than 90 filamin-binding partners have been identified, including intracellular signaling molecules, receptors, ion channels, transcription factors, and cytoskeletal and adhesion proteins (Razinia *et al*. 2012).

Surprisingly, defects in denticle numbers in *jbug^30^* embryos are rescued by expression of Jbug-<u>PI</u>, an isoform that contains long IG-FLMN repeats without the actin-binding domains, but only minimally by expression of Jbug-PF (Figure 5). These data suggest a major role of Jbug-PI in denticle formation (Figure 9). Although Jbug is upregulated in the anterior side of each segment in the embryo, where denticle belts form, Jbug is not enriched in the denticles themselves (Figure 4). Interestingly, another *Drosophila* filamin – Cher – is enriched in denticles, colocalizing with actin protrusions (Dilks and DiNardo 2010). The different subcellular localization of Jbug and Cher in denticle forming cells suggests distinct roles for the two *Drosophila* filamins. Cher likely acts as a component of denticles that organizes actin filaments. A simple model for Jbug function is that actin-binding domains might act to stabilize actin filaments and that protein-protein interactions through long IG-FLMN repeats might recruit other key components for denticle formation to the apical plasma membrane where actin-rich protrusions arise. Still, Jbug-RF/M/N and/or Jbug-RL appear to contribute to denticle formation. Although overexpression of *jbug-RF* in *jbug^20^* or *jbug^30^* mutant embryos does not fully rescue the decrease of denticle numbers (Figure 5), zygotic knockdown of *jbug-RF/M/N* and *jbug-RL* using the RNAi line specifically targeting them (*GD13033*) results in premature/small denticle formation and also decreases the number of denticles in some rows (Figure 5), suggesting roles for the actin-binding domain-containing isoforms in denticle formation (Figure 9).

### Jbug is likely to affect PCP indirectly

*jbug^30^* and *jbug^133^* alleles show a delay of actin-rich prehair formation in the pupal wing and a mild swirling pattern of adult wing hairs (Supplemental Figure 3). Both phenotypes are often shown in mutants of PCP genes (Adler, 2002). *jbug* mutants, however, do not show any defects in ommatidial rotation, suggesting that Jbug is less likely to directly affect the core PCP pathway. Rather, the phenotypes in pupal/adult wing hairs are likely to be caused by defects in formation of actin protrusions, similar to what we observe in thoracic bristles (Figure 2) and denticles (Figure 5) in *jbug* mutants. Consistent with this idea, Jbug has been proposed to modulate the mechanical properties of actin filaments in tendon cells to regulate planar orientation of thoracic bristles; alterations in the ability to adjust tensile forces upon *jbug* knockdown disrupt the planar orientation of bristles (Olguin et al., 2011).

### Roles of Jbug in formation of tubular organs

Several studies suggest roles for filamin A in tubular epithelial morphogenesis and diseases in tubular organs. During branching morphogenesis of mammary glands, the levels of filamin A expression and the extent of the integrin-filamin interaction modulate collagen remodeling to optimize matrix stiffness to support tubulogenesis (Gehler *et al*. 2009). Filamin A is also an important regulator for Polycystin-2, a key molecule in autosomal polycystic kidney disease (Wang *et al*. 2015). Our data also suggests a role for Jbug in apical domain reorganization in tubular epithelial cells (Figures 7 and 8). Jbug shows strong apical enrichment in the early embryonic trachea and the salivary gland (Figure 6) and both Jbug loss and overexpression cause defective tube morphology with disrupted Crb localization along the apical surface of salivary gland cells (Figures 7 and 8). These data support the idea that appropriate levels of Jbug are important for apical organization during tubular organ development. Loss of apical Jbug enrichment, however, does not cause overt defects in tubular organ formation in *jbug* null or hypomorphic alleles (Figure 6). In these alleles, abundant cytoplasmic Jbug signals still remain, and the levels or distribution of Crb was not noticeably changed (Supplemental Figure 5). Overexpressed Jbug also mainly localizes to the cytoplasm in salivary gland cells, with only slight apical enrichment (Figure 7). These data suggest that apical enrichment may not be necessary for Jbug function in the formation of tubular epithelia.

### Transvection or trans-splicing might occur in the *jbug* locus

The surprising rescue that we have observed of viability, bristle morphology and wing hair orientation of *jbug^30^* and to a lesser extent with *jbug^133^* with *Exel6079*, a deficiency that deletes all *jbug* exons upstream of the transcription start sites of *RC* and *RI* (Table 1; Supplemental Figure 3), suggests epigenetic regulation. One possible mechanism is transvection, an epigenetic phenomenon that results from interactions between alleles on homologous chromosomes in tissues where the chromosomes are paired (King *et al*. 2019), typically with enhancers of one allele on one chromosome affecting expression of the other allele on the homologous chromosome. Here, however, we have to propose transvection via template-switching, wherein transcripts from the *jbug^30^* allele would switch templates at some point outside the *Exel6079* deficiency but before the large insertion in *jbug^30^*, for example. Another possibility is trans-splicing, a gene regulatory mechanism that joins exons from two separate transcripts to produce chimeric mRNA, and has been detected in most eukaryotes (Horiuchi and Aigaki 2006; Lasda and Blumenthal 2011). In *Drosophila*, trans-splicing has been demonstrated to be an essential process for two genes, *longitudinals lacking* (*lola*) and *modifier of mdg4* (*mod(mdg4)*). In the *lola* locus, the 5’-end of *lola* transcripts contain five exons that splice into 20 variants of 3’ exons to generate 20 protein isoforms (Ohsako *et al*. 2003). In the *mod(mdg4)* locus, two mutant alleles of the gene, each located in one of the two homologous chromosomes, restore the wild-type function of the gene (Mongelard *et al*. 2002), much like is observed here. The presence of Jbug-PF/M/N isoforms (and the rescue of PI size) in transheterozygous animals for *jbug* hypomophs and *Exel6079* (Figure 1G) supports the idea that trans-splicing might occur, where the 5’ exons of *jbug-RF/M/N* transripts on *jbug^30^* or *jbug^133^* are spliced to 3’ exons of *jbug* transcripts on the *Exel6079* chromosome. Recent studies have revealed conserved sequences in the *mod(mdg4)* intron that promote trans-splicing in this gene (Gao *et al*. 2015; Tikhonov *et al*. 2018). Identification of the sequences that promote trans-splicing in the *jbug* locus will help better understand this surprising gene regulatory mechanism.

---

**Supplemental Figure 1.**
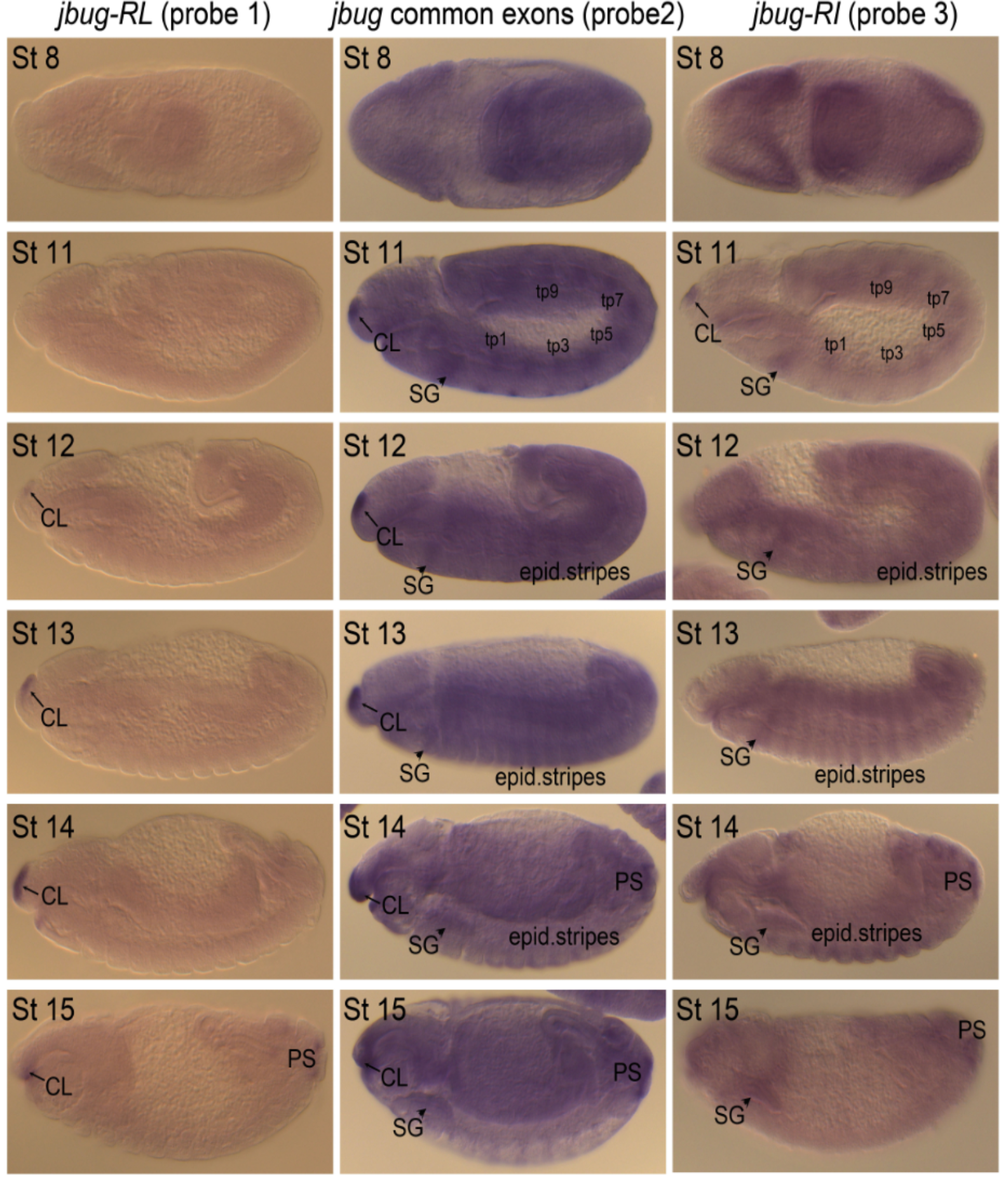
In situ hybridization. In situ hybridization of wild-type embryos using probes specific for different *jbug* splice forms as shown in Figure 1A. CL, clypeolabrum. SG, salivary gland. tp, tracheal pits. PS, posterior spiracles.

**Supplemental Figure 2.**
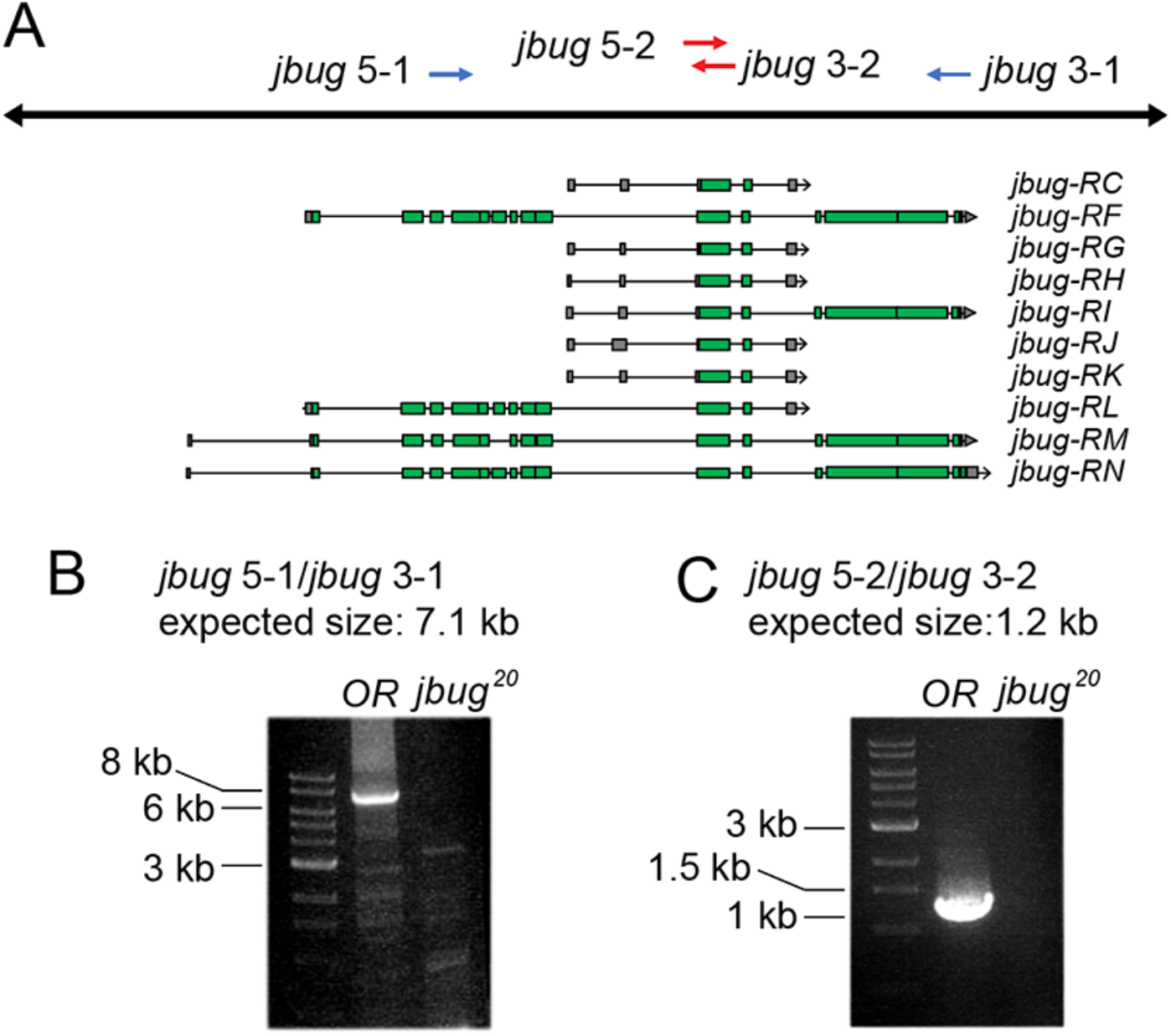
Reverse transcriptase-PCR (RT-PCR) reveals a loss of *jbug* mRNA in *jbug^20^* null mutants. (A) Genomic structures for the jbug locus. Blue and red arrows represent primers used for RT-PCR. (B and C) RT-PCR using cDNAs made from wild-type (OR) and *jbug^20^* homozygous embryos. Whereas RT-PCR using wild-type cDNA produces a PCR product of expected sizes (left lanes in B and C), no bands are detected in *jbug^20^* (right lanes in B and C).

**Supplemental Figure 3.**
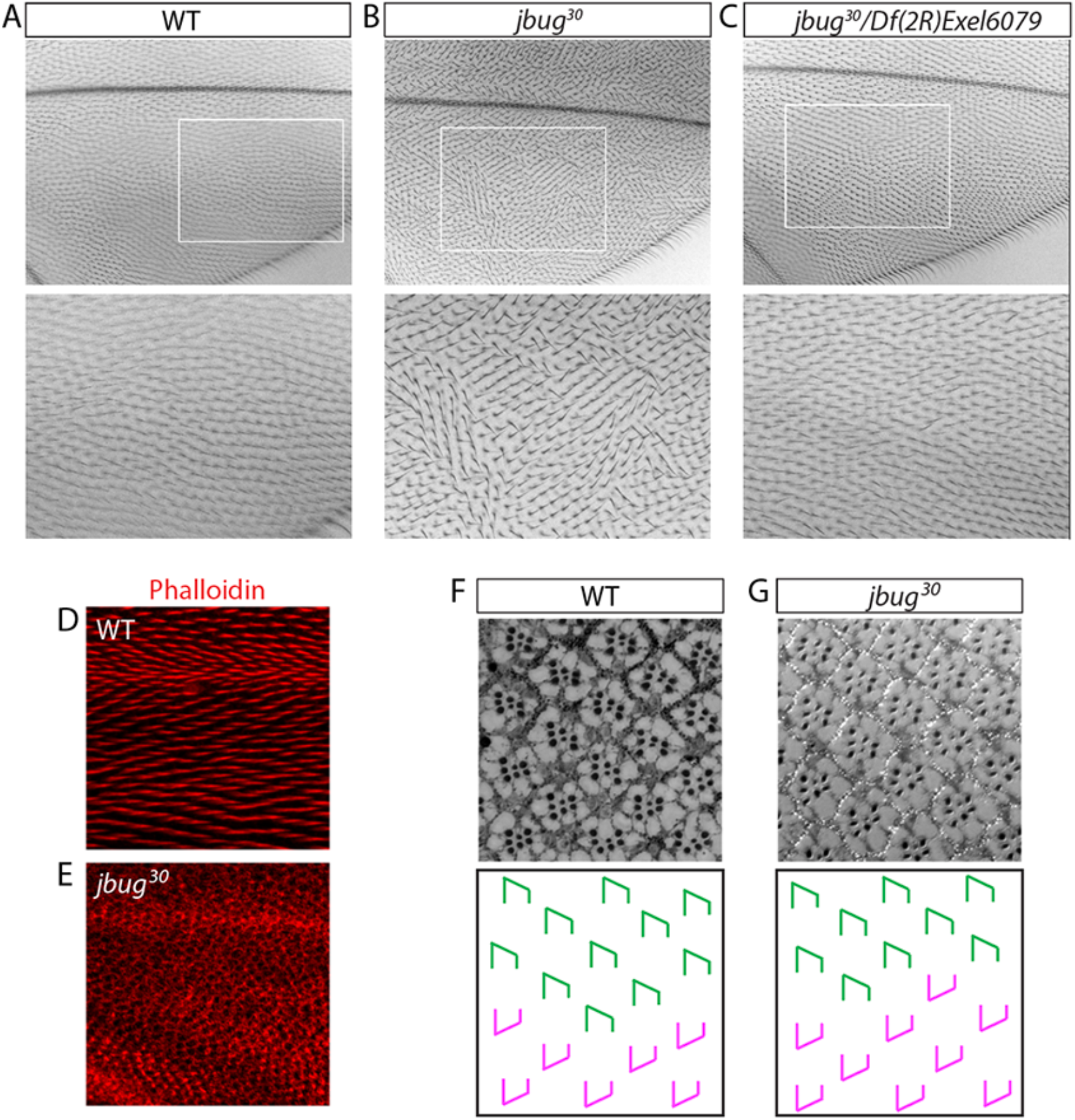
Mutations in *jbug* cause mild planar cell polarity defects in adult wing hair orientation. (A-C) Adult wings. Compared to the largely distal orientation of wing hairs in the wild-type wing (A), wings in *jbug^30^* mutants (B) show mild swirling patterns of wing hairs. (C) *jbug^30^/Exel6079* flies do not show defects in the wing hair orientation. White boxed regions are shown in higher magnification in the bottom. (D, E) Wild-type (D) and *jbug^30^* mutant (E) pupal wings (32 hours APF) stained for phalloidin. Prehair formation is delayed in the pupal wing in *jbug^30^*. (F, G) Adult ommatidia of wild type (F) and *jbug^30^* (G) near the dorsal/ventral boundary, the equator. Schematic drawings are shown in the panels below the actual images. Green and magenta shapes indicate the orientation of ommatidia normally found in the dorsal and ventral hemisphere of the eye, respectively. *jbug^30^* flies do not show planar cell polarity defects in the orientation of ommatidia.

**Supplemental Figure 4.**
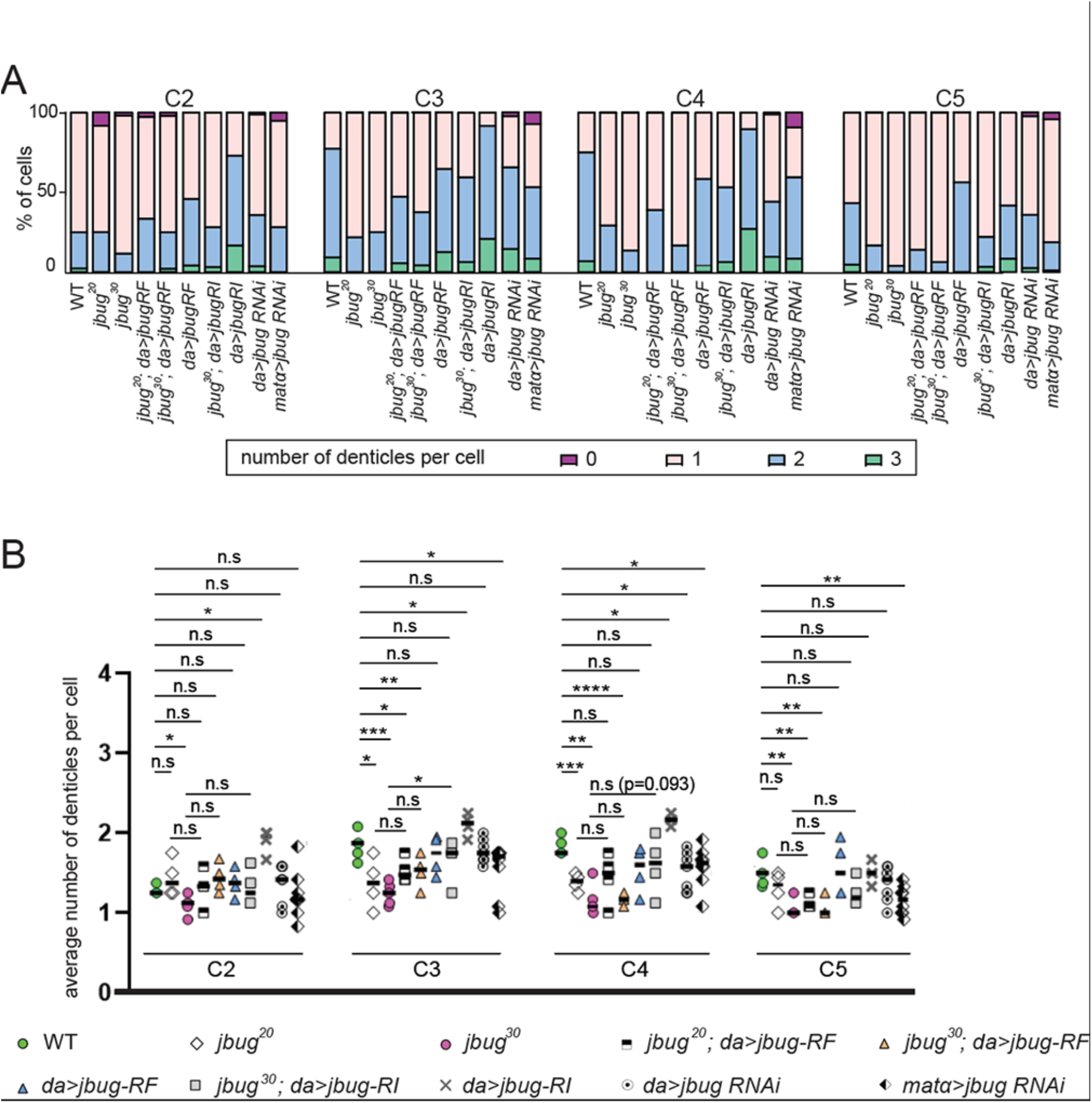
Jbug regulates the formation of denticle precursors in the ventral epidermis of the embryo. (A) Percentage of cells that have different numbers of denticles in rows C2-C5. (B) Quantification of the average number of denticles per cell in C2-C5. Student’s t-test with Welch’s correction is used for statistics calculation. The same four cells as shown in Figure 5K were chosen for quantification for each genotype.

**Supplemental Figure 5.**
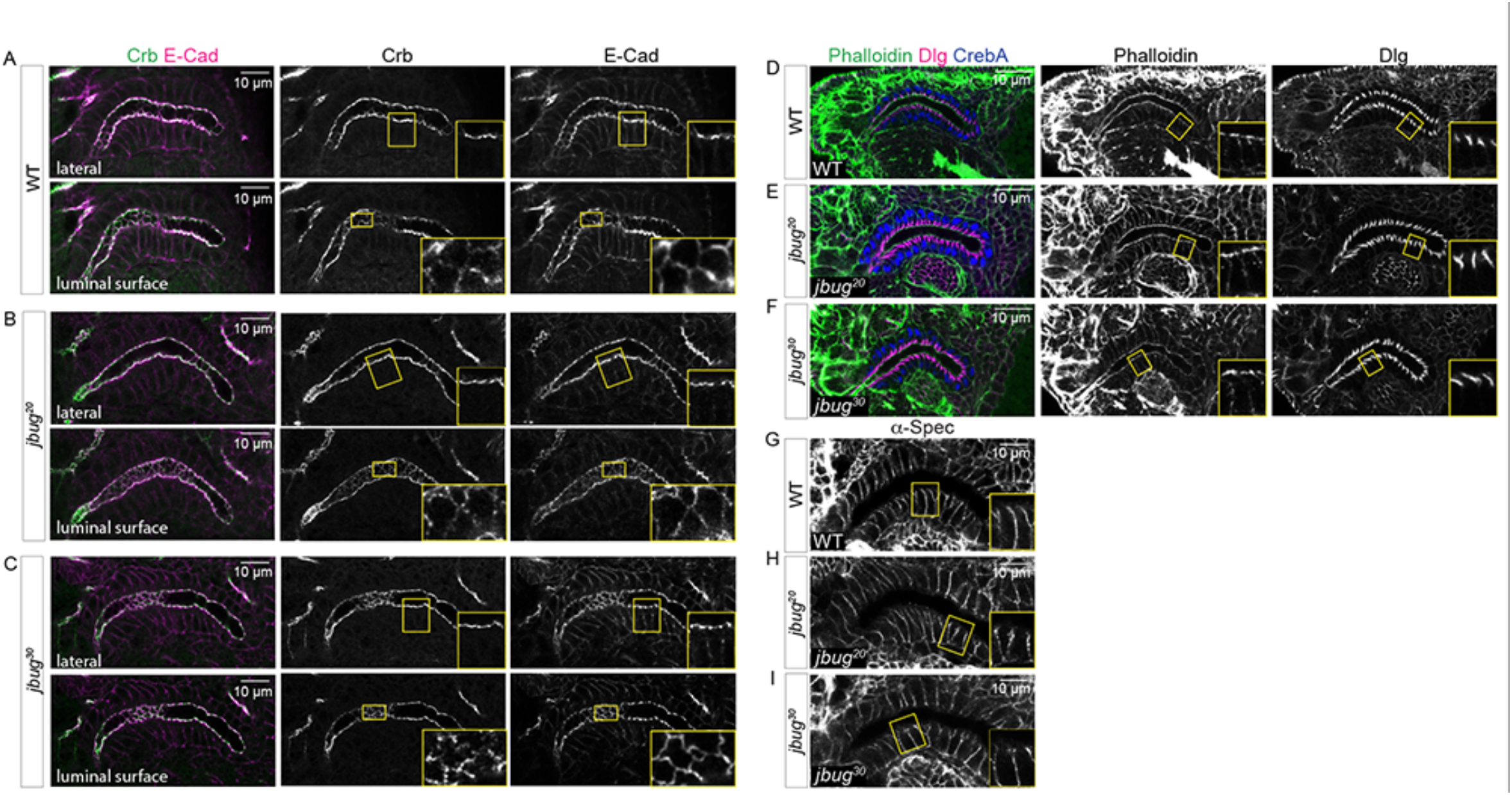
Epithelial polarity markers and F-actin and are not affected in salivary glands in *jbug* mutants. (A-I) Confocal images of stage 16 salivary glands. (A-C) Wild-type (A), *jbug^20^* (B) and *jbug^30^* (C) salivary glands stained for Crb (green) and E-Cad (magenta). (D-F) Wild-type (E), *jbug^20^* (F) and *jbug^30^* (G) salivary glands stained for phalloidin (green) and Dlg (magenta). (G-I) Wild-type (G), *jbug^20^* (H) and *jbug^30^* (I) salivary glands stained for α-Spec. Insets, higher magnification of the boxed region.

**Supplemental Figure 6.**
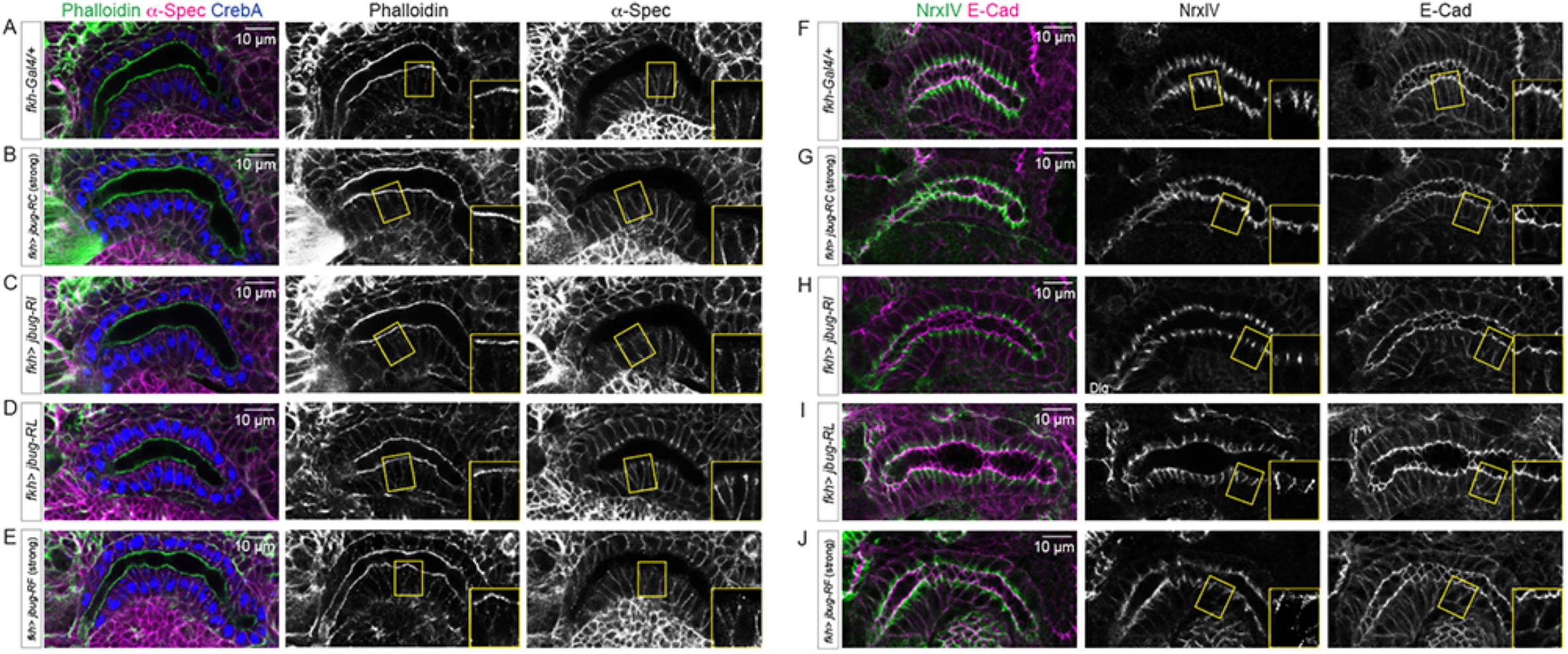
F-actin, epithelial polarity markers and junction markers are not significantly altered in salivary glands overexpressing different Jbug isoforms. (A-J) Confocal images of stage 16 control (*fkh-Gal4/+*, A and F) and salivary glands overexpressing Jbug-RC (B and G), Jbug-RI (C and H), Jbug-RL (D and I) and Jbug-RF (E and J). (A-E) Salivary gland stained for phalloidin (green), α-Spec (magenta) and a salivary gland nuclear marker CrebA (blue). (F-J) Salivary glands stained for Nrx-IV (green) and E-Cad (magenta). Insets, higher magnification of the boxed region.

**Supplemental Figure 7.**
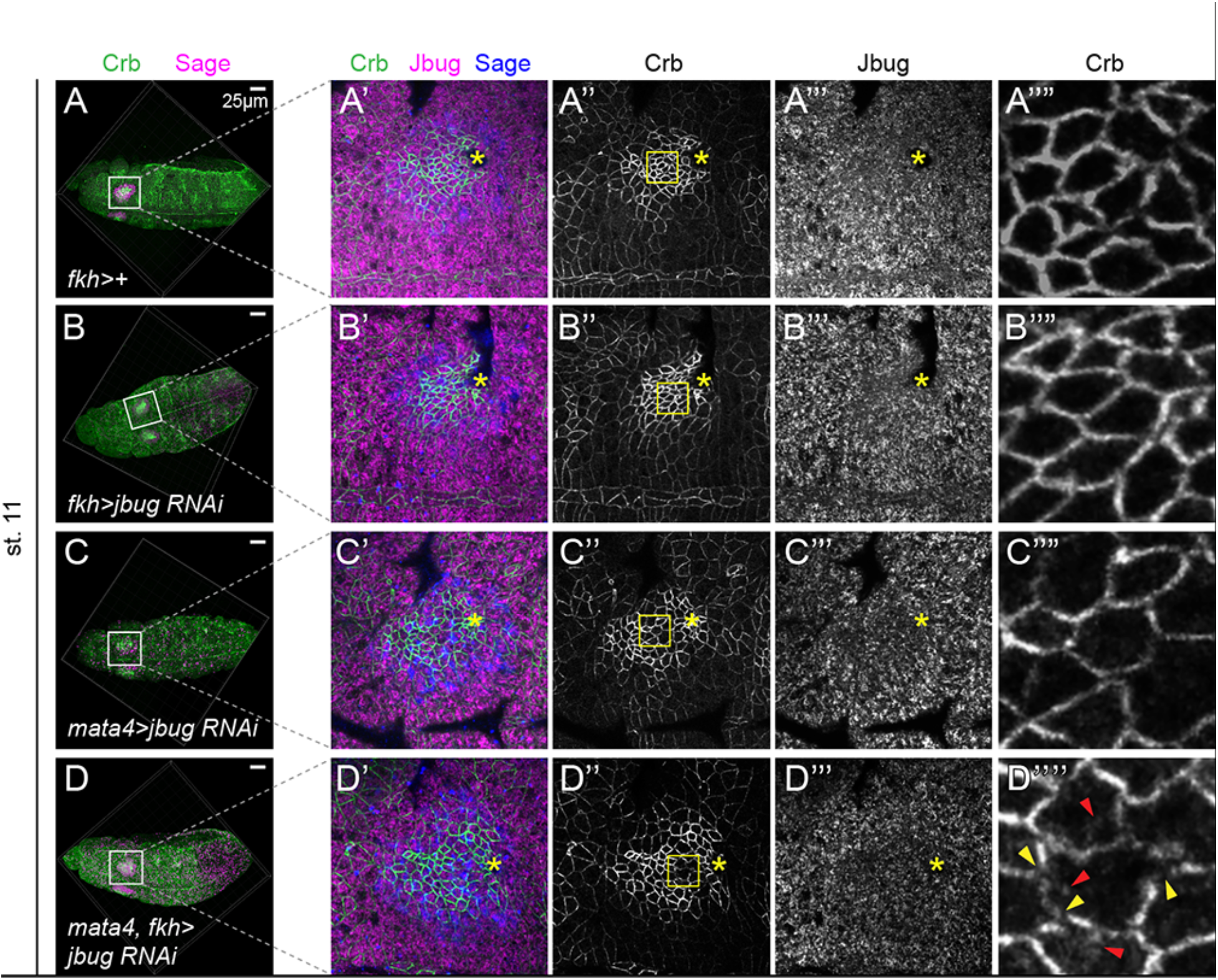
(A-D) Confocal images of stage 11 embryos stained for Crb (green), and Sage (blue). Control (*fkh-Gal4/+*; A), zygotic (*fkh>jbug RNAi*; B), maternal (*matα>jbug RNAi*; C), and both maternal and zygotic knockdown of *jbug* (*matα, fkh>jbug RNAi*; D). (A’-D’’’) Higher magnification of the salivary glands in A-D. Crb (green), Jbug (magenta) and Sage (blue) signals are shown. (A’’-D’’’) Crb (A’’-D’’) and Jbug (A’’’-D’’’) only. Jbug signals are slightly reduced in *jbug* knockdown, with a more significant reduction in maternal (C’’’) and maternal/zygotic knockdown (D’’’). (A’’’-D’’’’) Higher magnification of the boxed region in A’’-D’’. Maternal/zygotic knockdown of *jbug* results in gaps (yellow arrowheads) and mislocalized Crb signals to the apical surface (red arrowheads).

## Acknowledgments

We thank the members of the Chung and the Andrew laboratories for their comments and suggestions. We thank M. Mlodzik and the Bloomington stock center for fly stocks and the Developmental Studies Hybridoma Bank for antibodies. We thank Flybase for the gene information. This work is supported by start-up fund from Louisiana State University to S.C. and NIH R01 DE013899 to D.J.A.

## Notes

### Competing Interest Statement

The authors have declared no competing interest.

### Summary of Updates

Figures and the text have been updated with new data. Figures have been rearranged.

